# Biofilm competency of *Desulfovibrio vulgaris* Hildenborough facilitates colonization in the gut and represses adenoma development in a rat model of colon cancer

**DOI:** 10.1101/2022.07.26.501613

**Authors:** Susheel Bhanu Busi, Kara B. De León, Dan R. Montonye, Judy D. Wall, James Amos- Landgraf

**Author notes:** Corresponding author: James Amos-Landgraf. These authors contributed equally (Susheel Bhanu Busi, Kara B. De León). Systems Ecology Group, Luxembourg Center for Systems Biology, University of Luxembourg, 7, avenue des Hauts-Fourneaux, L-4362 Esch-sur-Alzette, Luxembourg (Susheel Bhanu Busi). University of Oklahoma Microbiology and Plant Biology Department, 770 Van Vleet Oval, Norman, OK 73019.

## Abstract

Human epidemiological and animal model studies have shown that the presence of colon cancer is associated with certain microbiota. Previous human colon cancer case-control reports and our preclinical model studies identify several bacterial taxa correlating with the suppression of tumor growth. This includes a sulfate-reducing bacteria from the genus *Desulfovibrio,* which correlates with fewer tumors and decreased phenotype penetrance as early as 1 month of age in rats. However, other studies have shown that *Desulfovibrio* spp. are decreased in relative abundance in healthy patient controls. To address this disparity we treated PIRC rats, a model of human familial colon cancer that harbored a complex gut microbiota, with biofilm-forming and biofilm-deficient strains of the sulfate-reducing bacterium *Desulfovibrio vulgaris* Hildenborough (DvH). We found that the biofilm-forming DvH strain could stably colonize the rat colon, with the bacteria detected even at 3.5 months post treatment. The biofilm-deficient DvH mutant only transiently colonized the rat colon and was no longer detected in fecal samples one-week post treatment. The colonic adenoma burden at four months of age was significantly reduced in rats colonized with the biofilm-forming DvH compared to those treated with the biofilm-deficient DvH. We found a differential shift in the endogenous gut microbiota (GM) structure over time, with a notable increase in known mucin degraders in the buffer-treated control and biofilm-deficient DvH treated groups. These latter groups of rats also showed a higher level of expression of *MUC2* in the colon, which encodes for the sulfonated mucin 2 expressed primarily in the gut and trachea. We also detected more dissolved sulfide in the feces from rats treated with either buffer or biofilm-deficient DvH compared to rats colonized with the biofilm-competent DvH. We found that the *in vitro* biofilm-forming capacity of DvH enabled *in vivo* engraftment of this bacterium within a complex GM population, altering the bacterial composition, and significantly reducing tumor burden in PIRC rats.

## Introduction

The exact etiology of colon cancer is complex and occurs through numerous acquisitions of combinations of genetic and epigenetic changes. Likely more than one combination that result in colon cancer involve risk factors including genetic predisposition and environmental factors (Mattar et al., 2011). Due to the high prevalence of colorectal cancer (CRC) in industrially developed countries, it is thought that environmental influences such as a Western-style diet comprised of increased consumption of meat, fats and total calories, coupled with longer life expectancies are factors for disease susceptibility (Bishehsari et al., 2014). Epidemiological studies have suggested that microbial dysbiosis in the gut together with bacterial biofilms are key factors for disease (Beaumont et al., 2016; Chung et al., 2018; Dejea et al., 2014; Sobhani et al., 2013; Zou et al., 2018). However, the mechanisms behind the role of the complex gut microbiota (GM) and how commensal bacteria contribute to adenomagenesis have only been worked out in a few bacteria such as *Bacteroides fragilis* (Chung et al., 2018)and *Fusobacterium nucleatum* (Kostic et al., 2013; Kostic et al., 2012) but the mechanisms driving susceptibility with the majority of bacteria associated with diseases are largely unknown.

In the human gastrointestinal (GI) tract the complex GM is composed of approximately 10^14^ commensal bacteria, many of which help in the metabolism of organic and inorganic compounds (Thursby and Juge, 2017). Recent studies comparing normal epithelial and tumor tissues by culture-independent, 16S ribosomal RNA (rRNA) gene or shotgun metagenomic next-generation sequencing (NGS) methods have shown differences in specific bacterial relative abundances (Arthur et al., 2012; Chen et al., 2015; Garrett et al., 2009; Kostic et al., 2013; Marchesi et al., 2011; Sears and Garrett, 2014; Sekirov et al., 2010; Zackular et al., 2013). Similar to these reports, our previous study assessing the role of the complex GM on colon cancer susceptibility in a preclinical model of familial adenomatous polyposis (PIRC rats) found that *Desulfovibrio* spp. was elevated in the low tumor group, where two rats out of 33 rats did not develop any colonic tumors (Ericsson et al., 2015). Contrarily, Zhu *et al*. (2014) demonstrated that *Desulfovibrio* along with other taxa were significantly enriched in a rat model of CRC (Zhu et al., 2014). A meta-analysis based on a metagenomics datasets with 156 samples similarly showed a statistically significant increase in the abundance of *Desulfovibrio* in CRC patients compared to healthy controls (Ai et al., 2019). While this phenomenon of increased *Desulfovibrio* associated with CRC has been reported by several studies (Chen et al., 2012; Dahmus et al., 2018), other studies have shown that this particular taxa may not be involved in disease progression (Balamurugan et al., 2008; Scanlan et al., 2009).

To determine the role of *Desulfovibrio* spp. in disease susceptibility, we treated PIRC rats with *Desulfovibrio vulgaris* Hildenborough (DvH). DvH is a Gram-negative, sulfate-reducing bacterium isolated from clay in 1946 (Postgate, 1979) and is not known to cause disease. It is often used as a model sulfate reducer in research studies and has been used for several industrial applications (De León et al., 2017) including radionuclide bioremediation of toxic environmental contaminants (Chakraborty et al., 2012) and wastewater treatment (Santegoeds et al., 1998). It is also known to form biofilms, adhering to surfaces with protein filaments (Clark et al., 2007). We previously reported on a spontaneously biofilm-deficient strain of DvH (DvH-MO) that has 12 single nucleotide polymorphisms (SNPs) in the genome (De León et al., 2017). One of these mutations is in the ABC transporter gene of the type 1 secretion system (T1SS) and was shown to be the cause of biofilm deficiency. Two large proteins were identified as T1SS cargo proteins and shown to be localized on the cell surface only when the T1SS is active. At least one of these large proteins, with either being sufficient, was required for biofilm formation on glass. A similar large T1SS exported protein (BapA) that is required for biofilm formation in *Salmonella enterica* (Latasa et al., 2005) and *Acinetobacter baumannii* is also required for adherence to eukaryotic cells and colonization (Berne et al., 2015; Brossard and Campagnari, 2012). We hypothesized that a deficiency in the T1SS function that impaired biofilm formation in DvH would lead to reduced colonization capacity compared to colonization of the PIRC rat gut by the biofilm-competent DvH. To test this, we compared the gut colonization of a fluorescent biofilm-forming DvH (JWT733) strain and a control strain lacking the ABC transporter of the T1SS which caused a deficiency in biofilm formation (JWT716; De León et al., 2017). We treated PIRC rats with the biofilm - competent and -deficient strains of DvH to determine the effect of colonization on adenoma burden. We found that DvH, a sediment bacterium, was capable of colonizing the rat colon, but only when expressing the capacity to form biofilms on glass *in vitro*. This colonization led to a significantly reduced adenoma burden in PIRC rats. This is the first report of DvH colonization in the rat colon with a complex gut microbiota. We also found that the GM communities in the rats treated with biofilm-forming DvH were modulated by the bacterial treatment, leading to a decrease in sulfide levels detected in the fecal samples along with a concomitant decrease in expression of mucin production and DNA damage genes. This study demonstrates the requirement of the type 1 secretion system in DvH in colonizing a preclinical rat model of human colon cancer. Although the effect of hydrogen sulfide leading to DNA damage has been hypothesized previously, our results are consistent with the potential involvement of this compound in colon cancer of *in vivo* models.

## Results

### Type 1 secretion system (T1SS) ABC transporter essential for DvH colonization of PIRC rats

We previously reported that a single nucleotide polymorphism in DVU1017 conferring an alanine to proline change in the encoded T1SS transmembrane, ATP binding protein is the cause for biofilm deficiency in DvH-MO (De León et al., 2017). We postulated that this ABC transporter gene of the T1SS, required for biofilm-competency on glass is also essential for bacterial colonization *in vivo*. To determine if the colonization potential of a previously identified low tumor group taxa (*Desulfovibrio* spp.) affected disease burden, we conducted a preliminary study where we gavaged two groups of male PIRC rats with 2 x 10^6^- 2 x 10^7^ DvH cells; one group received wildtype, biofilm-competent (WT) cells and the other received biofilm-deficient DvH-MO cells. We found that at 1-week post treatment, 100% (4 of 4) of the rats treated with WT had DvH colonization of the gut (determined in fecal samples) and this colonization was maintained through 4 months of age. In contrast, only 17% (1 of 6) of the animals treated had detectable levels of the DvH-MO strain after 1-week. At 4 months of age, none of the rats showed detectable levels of the DvH-MO strain shed in feces. PIRC rats treated with the wildtype strain had a reduced average tumor area compared to tumors found in DvH-MO treated rats (Supplementary Fig.1B). The former had only 13% of the tumors larger than 5 mm^2^ (Supplementary Fig.1C). In contrast, rats treated with the mutant DvH-MO strain had several tumors (∼35%) that were larger than 5 mm^2^ in average area (Supplementary Fig.1B, 1C and Supplementary Fig.2). Although the average colonic tumor number did not differ significantly, the average size did (Supplementary Fig.1A). It was not clear that biofilm capacity was the feature causing this difference in the treated rats because DvH-MO has 12 additional sequence differences compared to WT. Therefore, to confirm that the T1SS mutation impeded DvH colonization of the rat colon, an isogenic strain derived from WT with a single deletion in the DVU1017 ABC transporter gene and designated JWT716 (De León et al., 2017) was used as a biofilm-deficient strain in our second study. We also generated a fluorescent, T1SS-competent, WT DvH strain expressing *dTomato* (designated JWT733) for detection via colonoscopy.

In the follow-up study we treated PIRC rats of both sexes at days 14 and 15 of age with either JWT733 or JWT716 in PBS, or anaerobic PBS alone, (*i.e*. the biofilm -competent, biofilm-deficient strains and control treatment, respectively). We used quantitative PCR with strain-specific locked nucleic acid (Handy et al., 2013) probes designed to identify single nucleotide variants, to determine the colonization potential of the two DvH strains. One week after treatment with JWT716, we were not able to detect any DvH in fecal samples, consistent with the observations in our preliminary study with the DvH-MO strain. We detected JWT733 in 100% and 83% of the fecal samples at one-week post treatment and 4 months of age, respectively. We concurrently used colonoscopy to assess colonization in the colon of the PIRC rats treated with the fluorescent JWT733 strain starting at 2 months of age. PBS-treated, control group rats and JWT716-treated rats were devoid of any observable fluorescence during the colonoscopy with the appropriate filters. Contrarily, colonoscopy of JWT733-treated rats revealed observable levels of fluorescence. We found such detectable levels of fluorescence in the rat colon at all timepoints of the JWT733-treated animals (2-, 3-, and 4-months of age; Table 1 and Supplementary Fig.3). To determine if the biofilm-competent JWT733 was indeed forming biofilms in the colonic epithelium, we used fluorescent in-situ hybridization (FISH) with a custom probe. In spite of technical issues with the sectioning and staining methodology with certain animal tissues, we found that of the thirteen rats in the JWT733 group successfully sampled, 40% had detectable levels of the bacteria in the lumen of the histology sections examined (Supplementary Fig.4).

**Table 1.**
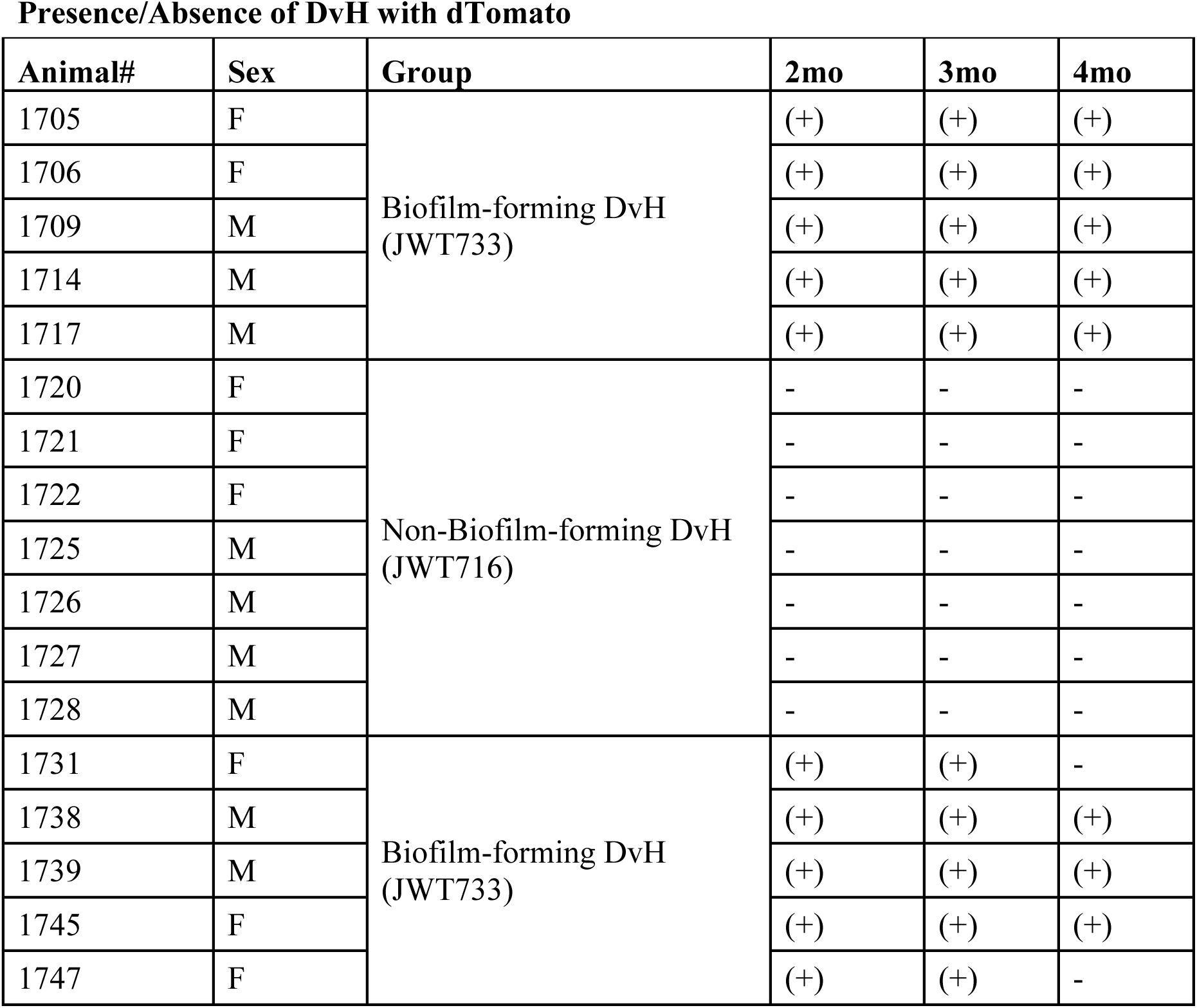
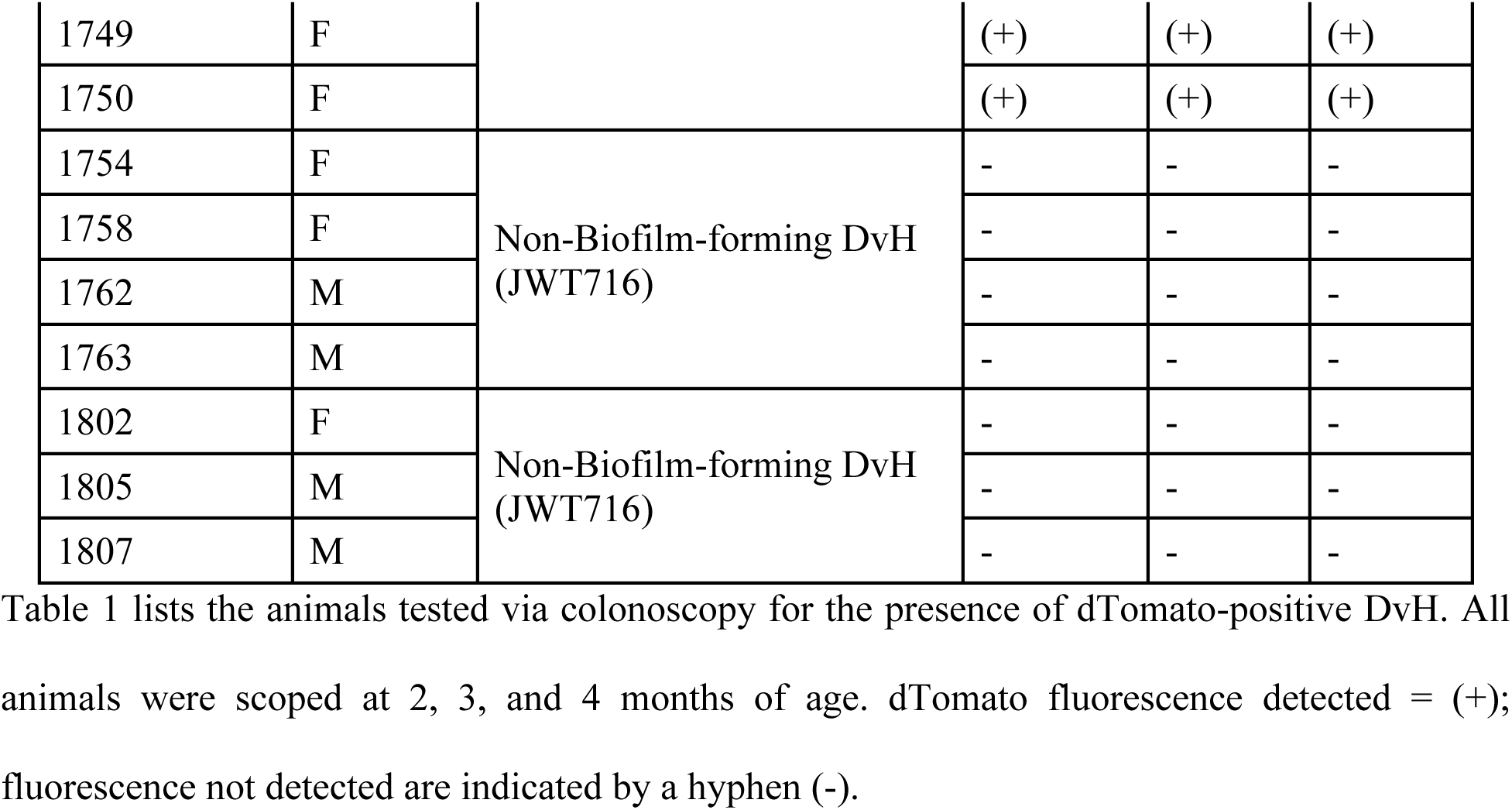
Colonoscopic assessment of dTomato presence or absence.

### Biofilm-competent DvH treatment associated with decreased adenoma burden

The biofilm-competent JWT733 treated PIRC rats, regardless of sex, had significantly reduced adenomas compared to the JWT716 (deficient in biofilm formation) or control groups (Fig.1A and 1B). The average size of the adenomas was also significantly reduced in the JWT733 group compared to the biofilm-deficient JWT716 group in the females, while the males showed a decreased average tumor area albeit statistically insignificant (Fig.1C and 1D). Here, we found that all tumors in the JWT733 group were smaller than or equal to 10 mm^2^ while the JWT716 and control groups respectively had 35% and 21% of tumors that were larger than 10 mm^2^ in size (Fig.1E). Using quantitative PCR, we determined the number of copies of the JWT733 bacterial strain from fecal samples at sacrifice and found that the colonic tumor number was inversely associated with the number of copies of JWT733 in the PIRC rats (Fig.1F).

**Figure 1.**
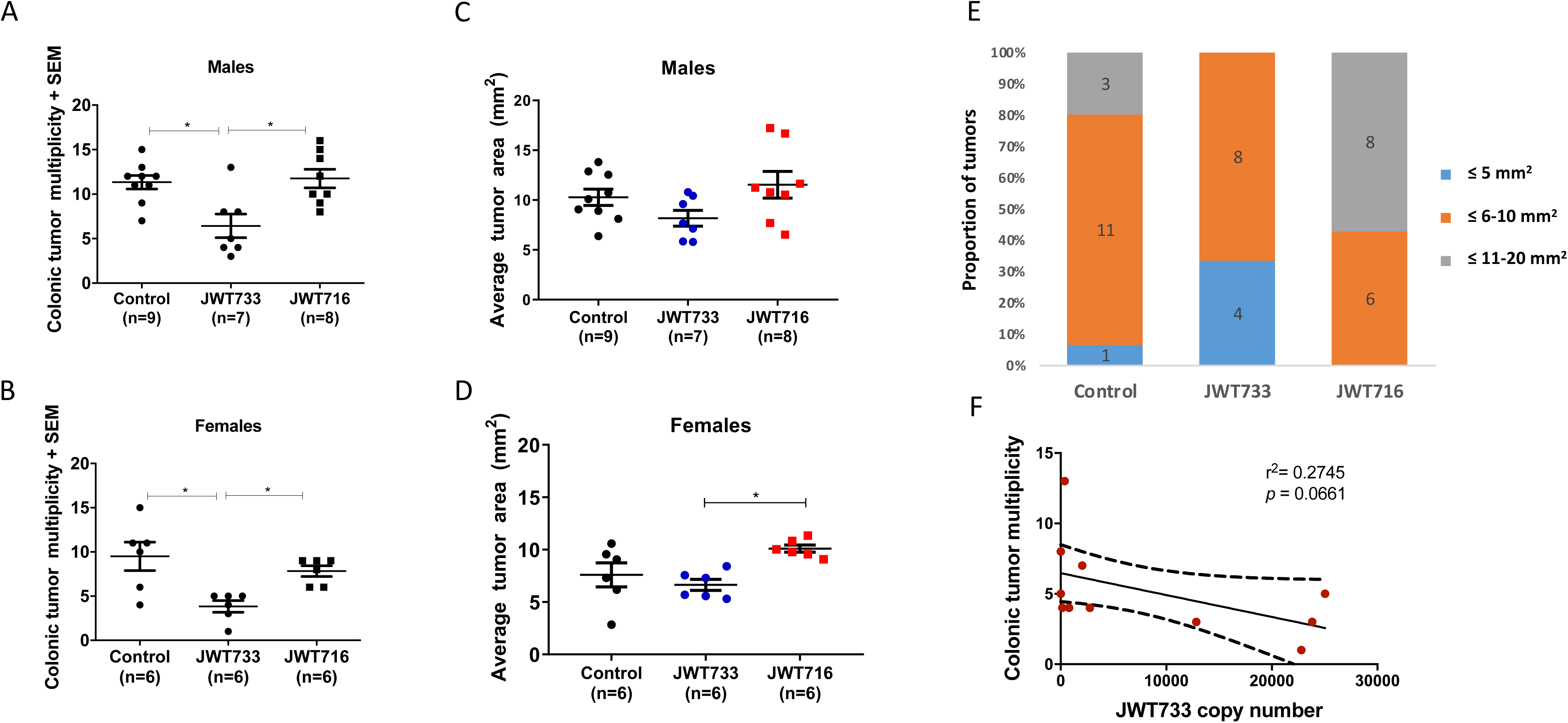
Tumor multiplicity, average tumor burden and OTU-tumor correlations in control and treated PIRC rats. Colonic tumor multiplicity in male (A) and female (B) PIRC rats at sacrifice, i.e. 4 months of age. Average tumor area observed in male (C) and female (D) PIRC rats treated with either anaerobic PBS (control), JWT733 (biofilm-competent) or JWT716 (biofilm-deficient) strains of DvH. For (A-D) a One-Way ANOVA with a Tukey’s post hoc test was used to determine significance with p-values below 0.05 considered to be significantly different between groups. (E) Tumor sizes observed in the treatment and control groups. Control, n=15; JWT733, n=12; JWT716, n=14. (F) Number of copies of JWT733 in DNA extracted from biopsies (collected at 2 months) of the biofilm-competent (wildtype) treated rats plotted against the colonic tumor multiplicity at 4 months of age. Error bars in all figures indicate standard error of the mean (±SEM). Dotted lines indicated the confidence intervals for the regression line.

### Fecal sulfide levels decreased in JWT733 treatment compared to the control and JWT716 groups

*Desulfovibrio* spp. is one of the many sulfate-reducing bacteria (SRB) found in the colon that serve as a source of sulfide in the GI tract (Rey et al., 2013; Tomasova et al., 2016). Based on the different capacities of gut colonization between DvH strains described above, we tested the concentrations of sulfide in the fecal samples. At necropsy (4 months of age), dissolved sulfide in fecal samples was not different between groups (Supplementary Fig.5A). However, at 2 months of age, we found that the high tumor groups -- JWT716 and control rats -- had significantly elevated levels of sulfide in the feces compared to the low tumor, biofilm-competent JWT733 treated group (Fig.2A). Due to the genotoxic nature of the hydrogen sulfide reported previously (Attene-Ramos et al., 2010), we examined the expression of DNA damage and repair genes in the normal colonic epithelium biopsies to assess the effect sulfide might be having on PIRC rats. We observed a significantly decreased expression (≥ 2 log2 fold-change) of the DNA damage repair genes such as *ATM*, *MSH2* and *MGMT* in the JWT733 group compared to the PBS-treated control rats or the biofilm-deficient JWT716 treated group (Fig. 2B). We also found higher expression of genes involved in hypoxia and inflammation in the colonic epithelium biopsies in the control and JWT716 rats compared to rats treated with JWT733 (Supplementary Fig.5B). The observation of higher expression of these inflammation-related genes is consistent with higher H2S levels. Overall, the genes associated with an increased inflammatory profile, *HIF1α* and *PTGS2* (Cox et al., 2004; Imtiyaz and Simon, 2010), demonstrated a significantly higher expression in the controls and JWT716 rats relative to the JWT733 treated rats.

**Figure 2.**
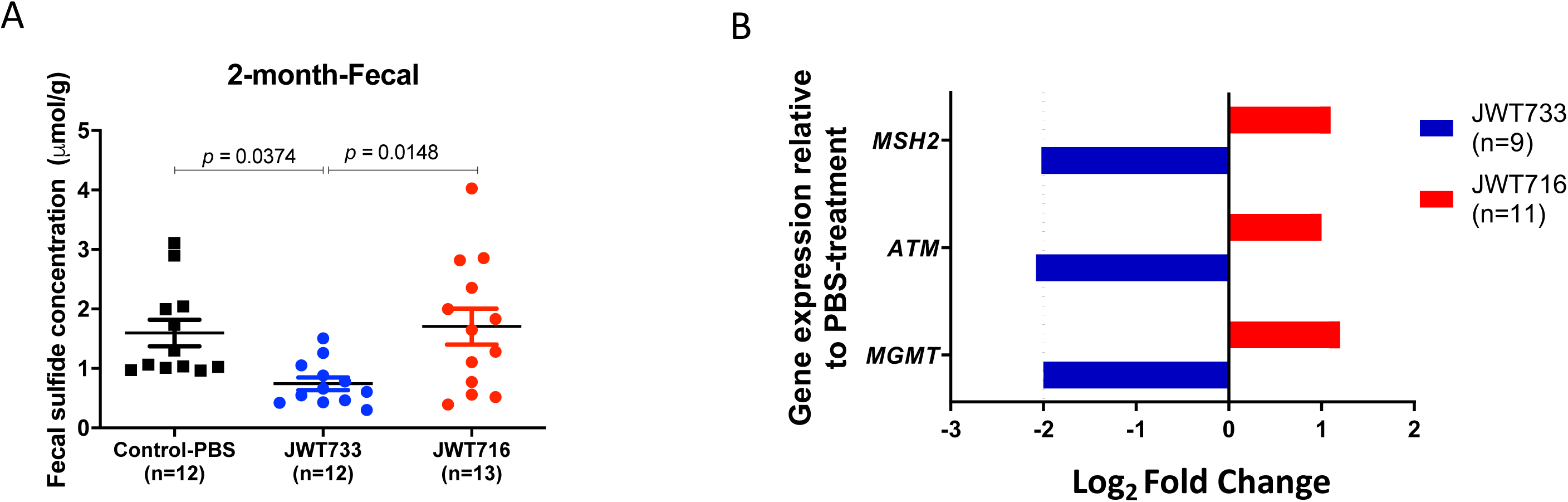
Sulfide assay and RT-qPCR gene expression of rat DNA damage genes. (A) Fecal sulfide concentration measured by Cline assay at 2 months of age in the control and treatment groups. p-values were calculated via a One-Way ANOVA with a Tukey’s post hoc test and those below 0.05 were considered to be significantly different between groups. (B) Relative expression of the DNA damage response genes relative to the PBS-treated control group (n=8) was determined by RT-qPCR. Log2 fold change was calculated using the ΔΔCq values, which is used to quantify the relative fold gene expression of samples. Red: expression in rats treated with JWT716 (biofilm-deficient, n=11); Blue: expression in rats treated with JWT733 (biofilm- competent, n=9) groups. Error bars in all figures indicate standard error of the mean (±SEM). Dotted line represents the 2-fold change between samples.

### DvH colonization modulates the complex GM architecture

Due to the different colonization capacities between the strains, we posited that the gut microbiota (GM) profile of the two groups would differ from each other post-treatment, thereby contributing to the different sulfide concentrations. We found that at one-week post-treatment there was a significant shift in overall profiles of the GM (Supplementary Fig.6A and Supplementary Table 3), which was observed even at 4 months of age between the wildtype- and mutant-treated groups (Two-Way ANOVA, Tukey’s post hoc, *p*<0.05; Supplementary Fig.6C).

We observed that the GM composition of the rats treated with the fluorescent, biofilm-competent strain (JWT733) and the parental, wildtype (WT) strain from the previous experiment were similar to each other based on the PERMANOVA analyses (Supplementary Fig. 7A and Supplementary Table 4). We found significant differences in the overall endogenous GM community structure between the control and each treatment group in the second experiment that were also different in sample types, i.e. fecal and biopsy (Fig.3A and Supplementary Table 5). Examination of all the significant OTUs (ANOVA, *p*<0.05) contributing to the differences in communities demonstrated different dominant OTUs in the fecal samples compared to the normal epithelium biopsy tissues (Fig.3B). We assessed differences at the phyla level in the fecal samples at 1-week post treatment (Fig.4A; Table 2). We found that the Proteobacteria phylum to which *Desulfovibrio* belongs, was significantly elevated in both the treated groups, compared to the control rats. While *Desulfovibrio* sequences were found in all three groups, the presence of *D. vulgaris* Hildenborough was only expected in the treated rats since it is not known to be part of the natural flora of the rat gut. To test whether *D. vulgaris* Hildenborough was present in all groups, we looked for the exact sequence of the 16S rRNA gene within the sequencing datasets. The *D. vulgaris* Hildenborough genome contains five 16S rRNA genes (identified as rrsA through rrsE) with four different sequences when analyzed across the full length (rrsC and rrsD are identical). For the region targeted for Illumina sequencing in this study, there are two possible sequences for *D. vulgaris* Hildenborough (that of rrsACDE and that of rrsB). The primer sequences were trimmed from raw Illumina short reads and the forward sequences were mapped to the *D. vulgaris* Hildenborough rrsA and rrsB sequences with Bowtie 2 and with default parameters (Langmead and Salzberg, 2012). Sequences that were identical to the reference sequences were counted within the resulting SAM file by identifying lines containing the SAM format predefined tag called “NM” of 0. This tag gives the number of differences (mismatches, insertions, and/or deletions) between the sequence and the reference. *D. vulgaris* Hildenborough was detected in JWT733-treated (n = 5 one-week samples at 0.002-0.58%) or JWT716-treated (n = 4 one-week samples and n = 2 two month biopsy samples at 0.002-0.09% relative abundance) rats. None of the control samples had sequences matching *D. vulgaris* Hildenborough. Thus, the *Desulfovibrio* spp. detected in the control rats were not *D. vulgaris* Hildenborough.

**Figure 3.**
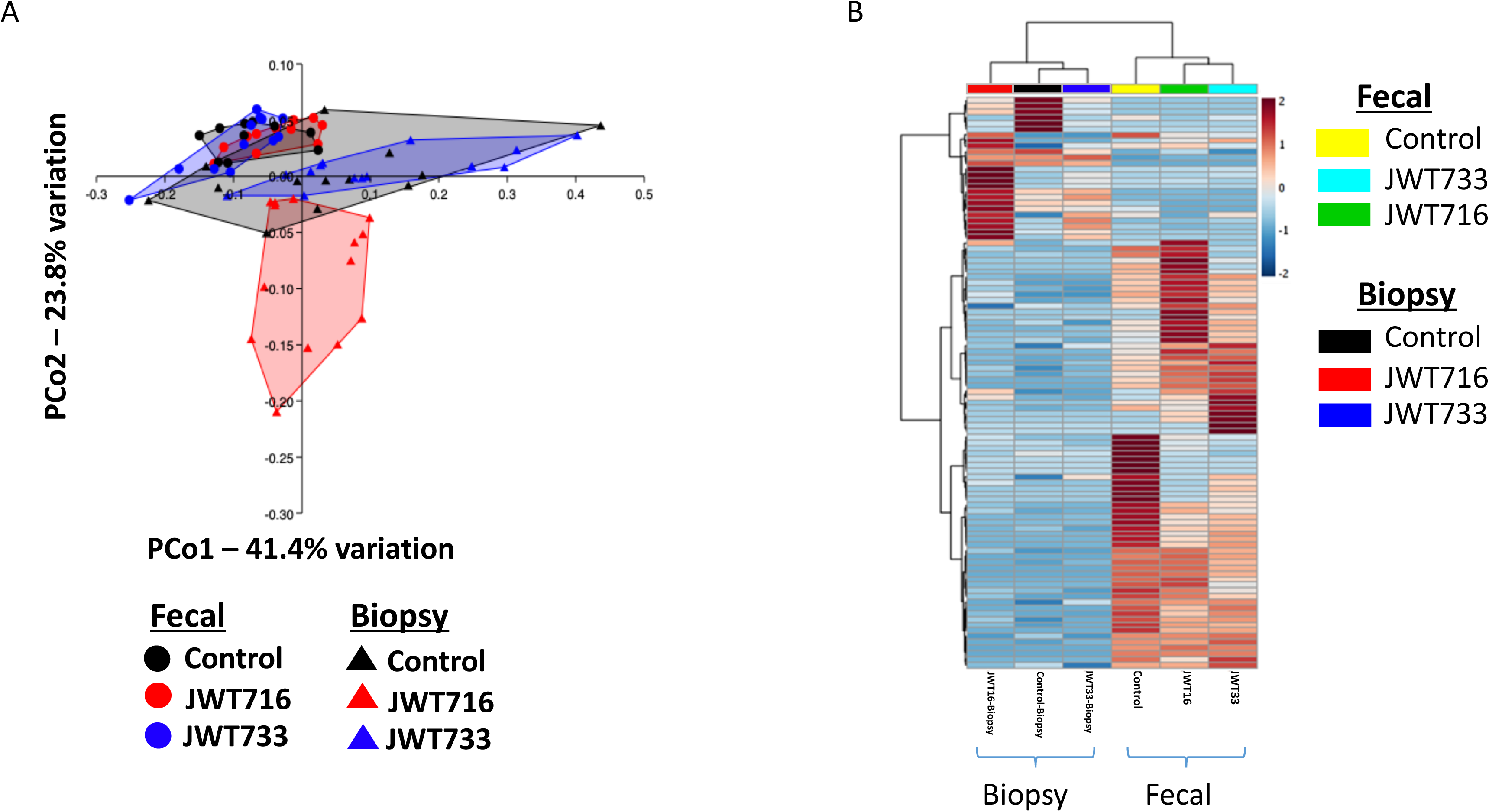
16S rRNA gene sequencing analysis of rats treated with JWT733, JWT716, or buffer. (A) Principal Component Analysis (PCoA) plot depicting the 16S rRNA gene sequencing dissimilarities between the groups at 2-months of age based on the Bray-Curtis distance matrix. Fecal samples are depicted as circles, while biopsy samples are shown as triangles. PBS: black, JWT733: blue and JWT716: green. Post-hoc analysis indicating the differences between individual groups is listed under Supplementary Table 5. Each symbol represents the GM community from the fecal sample of a single rat at 2 months of age. (B) Heatmap generated from the significantly (ANOVA, *p*<0.05) different OTUs between each group of the fecal and biopsy samples at 2 months, using Ward’s clustering algorithm. Range of blue to red color indicates low to high abundance respectively. PBS control, n=15; JWT733, n=13; JWT716, n=14 for both fecal and biopsy samples.

**Figure 4.**
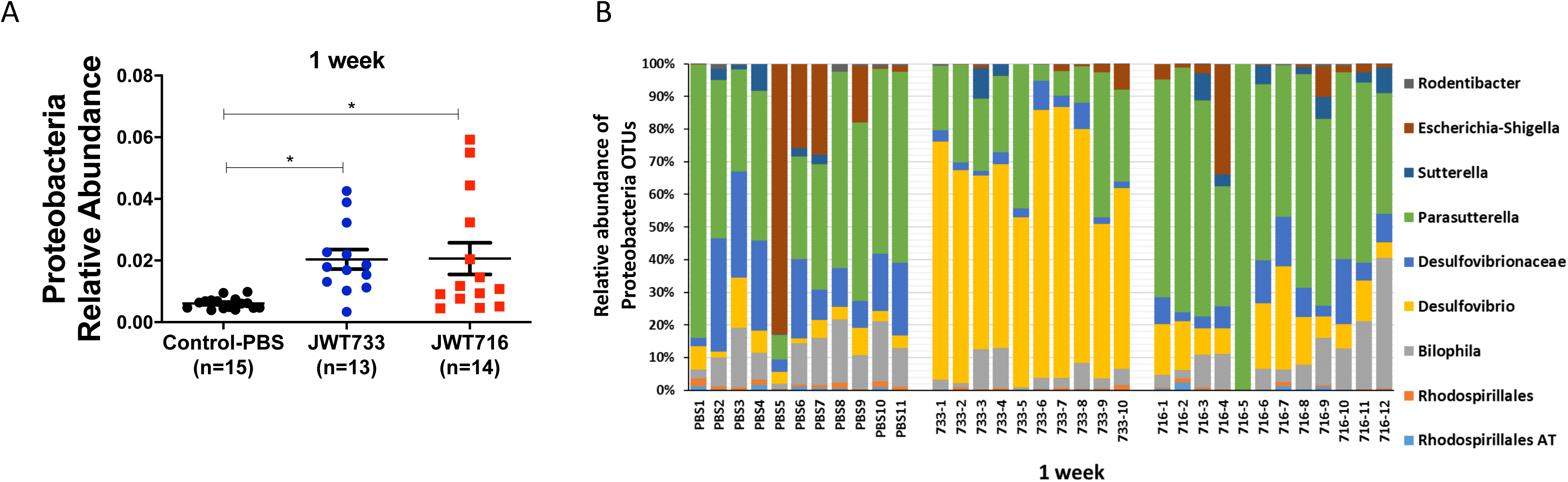
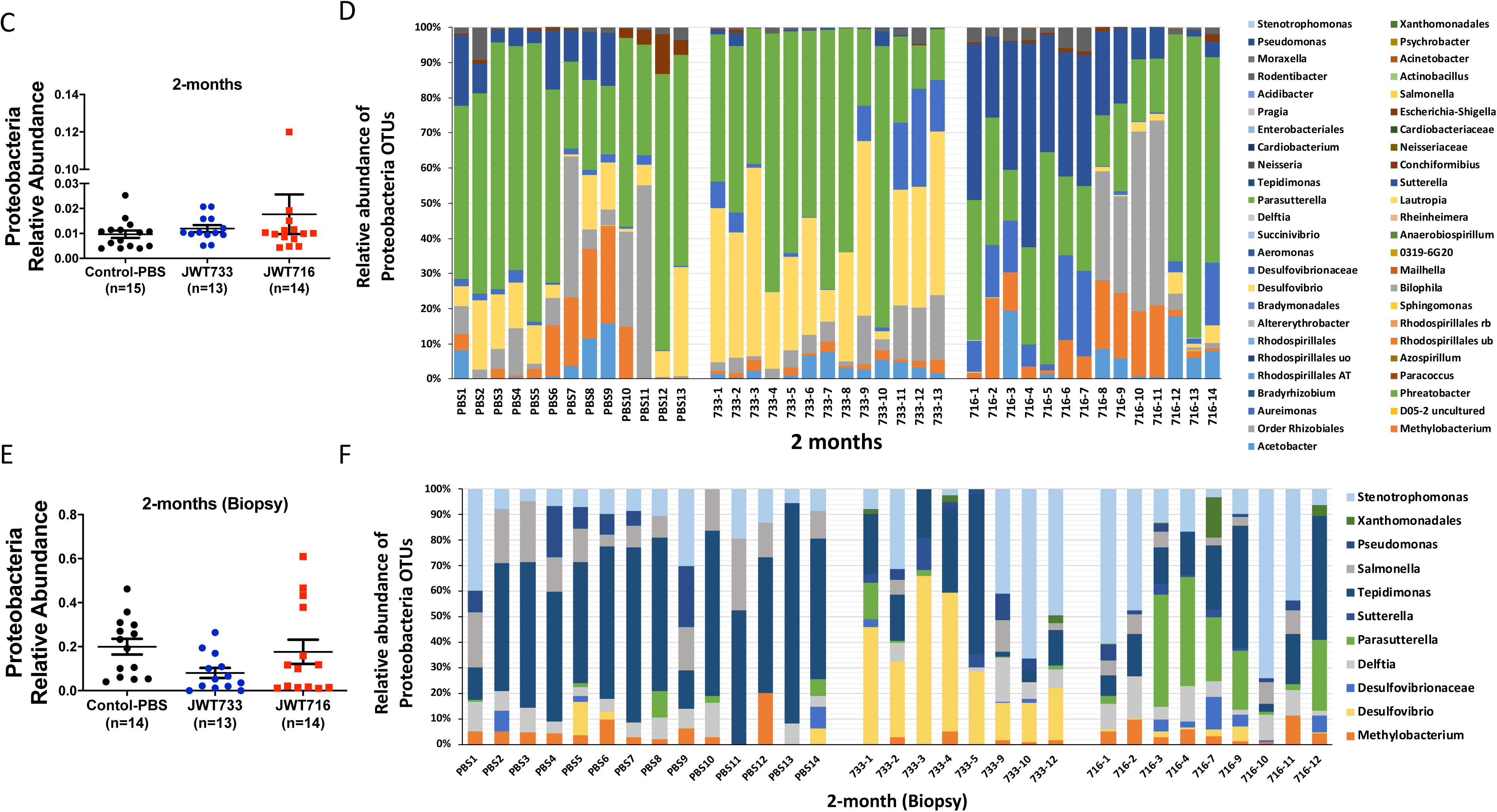
Proteobacteria species levels vary due to DvH-treatment. Dot plots and bar graphs depict the relative abundance of Proteobacteria (A) and the composition of the Phylum (B) respectively, in fecal samples at 1-week post treatment; 2-months of age (C) and (D), and mucosal biopsies at 2-months of age (E) and (F). Error bars indicate standard error of the mean (±SEM). Associated bar graphs show the relative abundance of the operational taxonomic units at the Genus level contributing to the Proteobacteria phylum in each sample. AT:ambiguous taxa; ub: uncultured bacterium, uo: uncultured organism and rb: rumen bacterium. Y-axes for figures (A), (C), and (E) represent the relative abundance percentage of the phylum and are to be observed as different at each time point.

**Table 2.** Proteobacteria changes in PIRC rat following isogenic *Desulfovibrio vulgaris* treatment

| DvH strain/<br>Treatment | Relevant Phenotype | Fecal Sample (following treatment) |  |
| --- | --- | --- | --- |
|  |  | One week | Two months |
| **JWT733** | Fluorescent Biofilm-forming (Lower tumor burden) | ↑<sup>A</sup> Proteobacteria vs Control | Proteobacteria same abundance (all treatments) ↑ *Desulfovibrio* vs Control<br>↑ *Desulfovibrio* vs JWT716 |
| **JWT716** | Biofilm-deficient (Higher tumor burden) | ↑ Proteobacteria vs Control<br>↑ *Parasutterella* vs JWT733<br>↑ *Escherichia-Shigella* vs JWT733 | Proteobacteria same abundance (all treatments) ↑ *Bilophila* vs JWT733<br>↑ *Methylobacterium* vs JWT733 |
| **Control (PBS)** | Buffer treatment | ↑ *Parasutterella* vs JWT733<br>↑ *Escherichia-Shigella* vs JWT733 | Proteobacteria same abundance (all treatments) ↑ *Bilophila* vs JWT733<br>↑ *Methylobacterium* vs JWT733 |

|  | (Higher tumor burden) |  |  |
| --- | --- | --- | --- |
| DvH strain/<br>Treatment | Relevant Phenotype | Biopsy Sample (following treatment) |  |
|  |  | One week | Two months |
| **JWT733** | Fluorescent Biofilm-forming (Lower tumor burden) | N/A | ↑ *Desulfovibrio* vs Control and JWT716<br>↑ *Stenotrophomonas* vs Control |
| **JWT716** | Biofilm-deficient (Higher tumor burden) | N/A | ↑ *Parasutterella* vs Control and JWT733<br>↑ *Stenotrophomonas* vs Control |
| **Control (PBS)** | Buffer treatment (Higher tumor burden) | N/A | ↑ *Pseudomonas* vs JWT716 and JWT733<br>↑ *Salmonella* vs JWT716 and JWT733 |

<sup>A</sup>Up-arrow is increased abundance; down-arrow is decreased. Relative changes in OTUs belong to the phylum Proteobacteria after treatment of PIRC rats with isogenic *Desulfovibrio vulgaris* Hildenborough strains and PBS are shown in Table 2. Changes are indicated for both fecal and biopsy samples at one week and at 2 months post-treatment. N/A: not applicable since biopsy samples were not collected at one week after treatment.

Additionally, at one-week post-treatment and at 2 months of age, we found variable levels of differential taxa under phylum Proteobacteria owing to JWT733 colonization or lack thereof of the JWT716 strain (Table 2; Fig. 4A and 4C). For example, at 1-week, *Parasutterella* and *Escherichia-Shigella* were enriched in both the control and non-biofilm-forming DvH group compared to the biofilm-forming group (Fig.4B). Simultaneously, at 2 months of age the overall relative abundance of Proteobacteria in the fecal (Fig.4C) and normal epithelium biopsy samples of the biofilm-forming group (Fig.4E) did not show any significant differences. Also, at 2 months, the differences observed at 1-week were replaced by the enrichment of *Bilophila* in the high-tumor groups (control and non-biofilm-forming DvH), along with *Methylobacterium* (Fig.4D) in fecal samples compared to JWT733-treated rats. Interestingly, *Parasutterella* was found to be abundant in the biopsy tissues of the non-biofilm-forming DvH-treated rats; whereas control rats showed elevated levels of *Pseudomonas* and *Salmonella* with a concurrent decrease in *Stenotrophomonas* when compared to the rats treated with JWT733 or JWT716 (Fig.4F).

Further examination of all the OTUs contributing to the fecal (Fig.5A) and biopsy (Fig.5B) GM profile differences among the three groups at 2 months of age identified different relative abundances of several OTUs (Table 2). In the fecal samples, we noticed taxa that were previously associated with an increased tumor phenotype (Flemer et al., 2017; Peters et al., 2016; Yang and Yu, 2018; Zhu et al., 2014) such as *Ruminoclostridium, Acetitomaculum, Bacteroidales,* and *Ruminococcaceae* V9D2013 were elevated in the control (PBS-treated) rats; whereas, in the non-biofilm forming strain-treated rats, we found increased abundance of *Fusobacterium*, and *Coprococcus* in fecal samples. In the rats treated with biofilm-competent JWT733, we found an increase in the abundance of taxa associated with decreased disease phenotype (Flemer et al., 2017; Yang and Yu, 2018; Zhu et al., 2014) including *Desulfovibrio, Bifidobacterium, Alistipes, Butyricimonas, Coprococcus, Erysipelotrichaceae.* Biopsies from the control rats, had elevated levels of *Tepidimonas* and *Anaerovorax*. In the JWT733 rats, we found similar increases of *Anaerovorax*, and *Roseburia* along with increased levels of bacteria such as *Bifidobacterium*, *Desulfovibrio*, *Eubacterium* and *Alistipes*. In contrast, biopsies of non-biofilm-forming JWT716 treated rats showed an enrichment in *Lachnospiraceae* UCG-008, *Methylobacterium*, *Staphylococccus*, and *Salmonella*.

**Figure 5.**
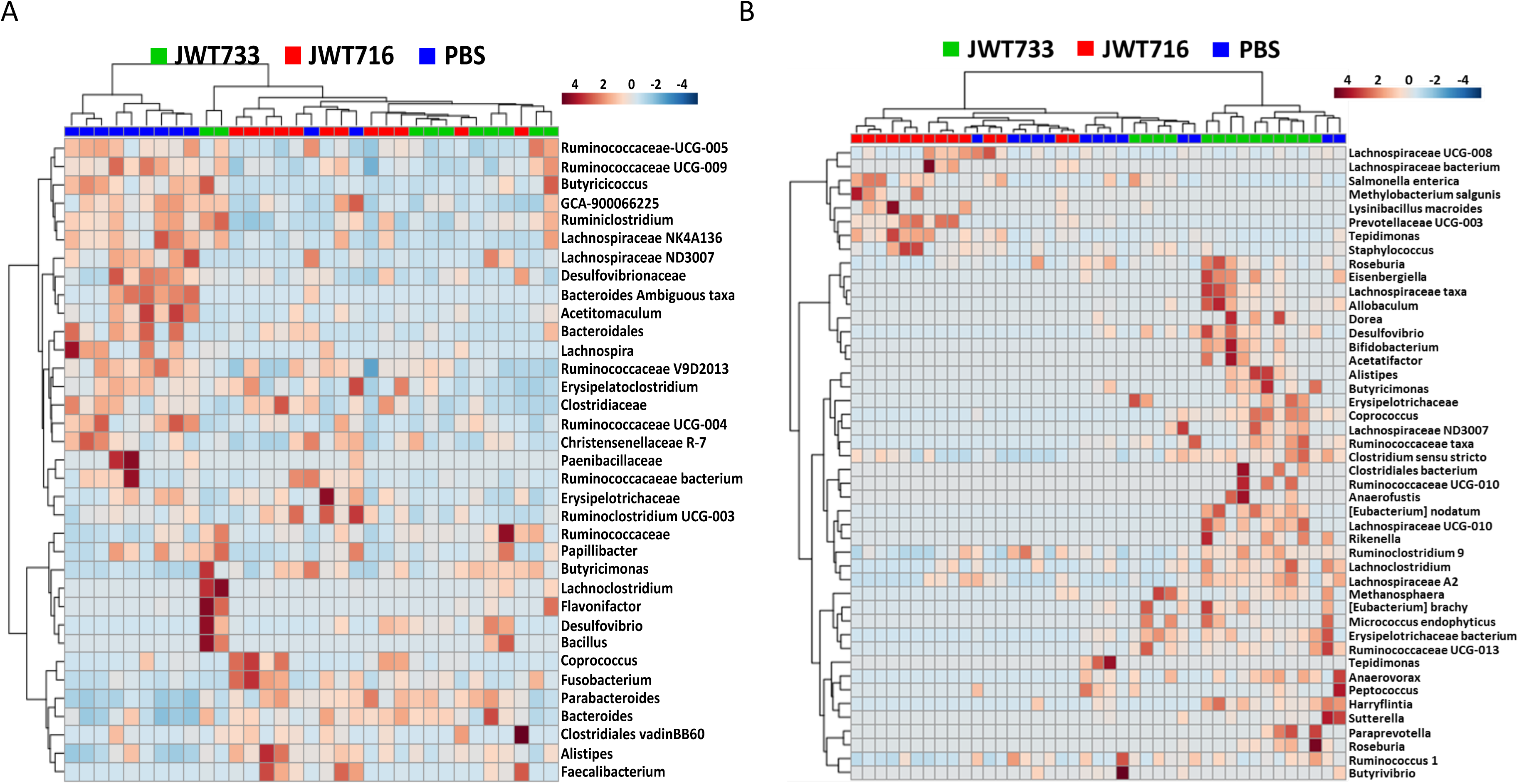
16S rRNA gene sequencing analysis of fecal and biopsy samples. (A) Heatmap of the GM profiles obtained from fecal (A) samples collected, or biopsy (B) samples collected at 2 months of age via colonoscopy. The OTUs shown are those that are significantly different (*p* < 0.05, DESeq2 analyses) between rat groups, clustered according to the Ward’s clustering algorithm. Range of blue to red color indicates low to high abundance respectively, generated based on Z-scoring, i.e. determining the standard deviation of sample from the mean of all samples for a particular taxon (generated sequentially for all OTUs). PBS, n=15; JWT733, n=13, JWT716, n=14, Con-biopsy, n=15; JWT733-biopsy, n=13 and JWT716-biopsy, n=14.

### Mucin-degrading bacteria associated with *MUC2* expression

Based on 16S rRNA sequencing with DESeq2 analyses accounting for false-discovery rates, we found significantly higher relative abundance in the fecal samples in mucin-degrading bacteria such as *Ruminococcaceae, Lachnospiraceae, Anaerotruncus, Bacteroides,* and *Mucispirillum* in the rat groups with higher tumor counts (i.e. those treated with PBS or JWT716; Fig.6A). However, we did not find similar differences in mucin-degrading bacteria in the biopsy samples across all three groups. It has been reported that mucin in the GI tract is an efficient source of sulfide (Carbonero et al., 2012). Mucin composition has been reported to be modulated by endogenous gut microbiota, where genera such as *Lactobacillus* and *Bacteroides* led to an increase in mucin gene expression. This has also been associated with an increase in abundance of mucin-foraging taxa such as *Ruminococcus*, *Bacteroides*, *Akkermansia*, and *Campylobacter* (Sicard et al., 2017).To determine if the increase in mucin-degraders correlated with increased mucus production, thus contributing to the H2S concentrations measured (Fig.2), we determined the expression levels of the primary rat gene encoding for mucin in the gut, i.e. *MUC2*. In biopsy samples, we found that at 2 months of age, *MUC2* expression was considerably lower in the rats colonized with JWT733 and had lower tumor counts compared to the rats in the high tumor groups (Fig.6B). We did not find any significant correlations between the *MUC2* gene expression and the relative abundance of the OTUs from fecal samples at one-week post treatment or 2 months of age; however, several OTUs did correlate with the colonic tumor number, potentially identifying predictive biomarkers of the disease (Fig. 6C and Supplementary Fig.8). We found a positive correlation with a mucin-degrader, *Ruminococcaceae* (Supplementary Fig.9A). Additionally, *Lactobacillus* and *Alistipes* negatively correlated with tumor counts (Supplementary Fig.9A and 9B).

**Figure 6.**
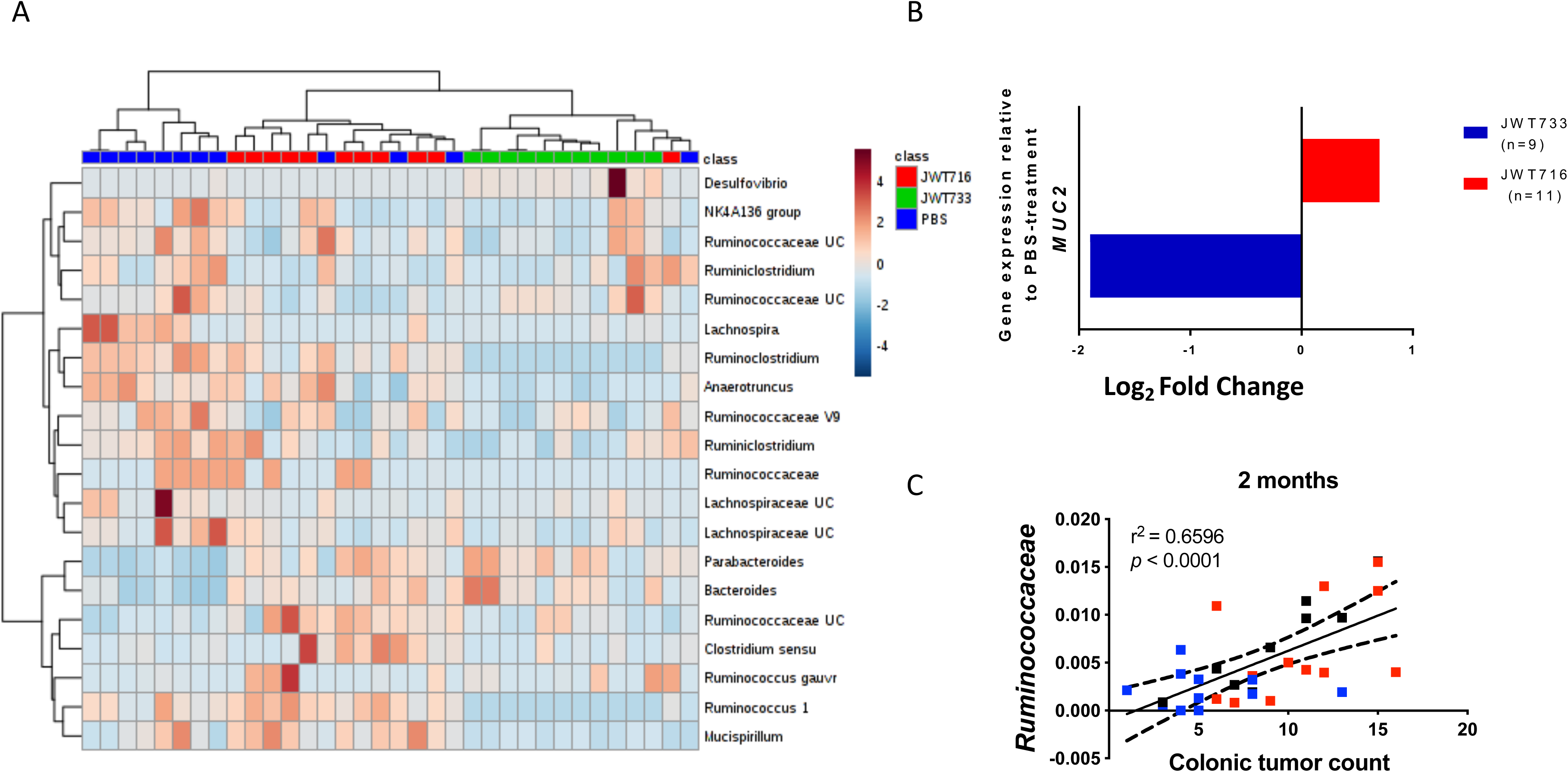
Mucin-degrading bacteria levels. (A) Heatmap of the mucin-degrading bacteria relative abundances obtained from fecal samples collected at 2 months of age depicting the top 25 significantly different OTUs between groups, using Ward’s clustering algorithm. Range of blue to red color indicates low to high abundance respectively, generated based on Z-scoring, i.e. determining the standard deviation of sample from the mean of all samples for each taxon. PBS, n=12, JWT733, n=11, JWT716, n=11. B) Relative expression of the gene involved in mucin-production relative to the PBS-treated control group (n=8) was determined by RT-qPCR. Log2 fold change was calculated using the ΔΔCq values, which is used to quantify the relative fold gene expression of samples. Red: expression in rats treated with JWT716 (biofilm-deficient, n=11); Blue: expression in rats treated with JWT733 (biofilm-competent, n=9) groups. C) Pearson’s correlations (*p*<0.05) between OTUs at one-week post-treatment with colonic tumor counts. Representative example of *Ruminococcaceae* (positive correlation) with colonic tumor count along x-axis and relative abundance of the taxa along then x-axis and relative abundance of the taxa along the y-axis is shown.

In summary, these data show that the ABC transporter gene in a type 1 secretion system of *Desulfovibrio vulgaris* Hildenborough, required for biofilm formation, is essential for colonization of the colon in PIRC rats. The colonization of biofilm-competent DvH in the rat colon was shown to be reproducible. Whether directly or indirectly, the T1SS of the bacterium is involved in reduced adenoma burden in this model of early onset colon cancer in the rat. We show that treatment with a bacterium in a complex GM setting could lead to significant shifts in the microbial community structure and affect host gene expression during this process. This is also a first report demonstrating that increased dissolved sulfide levels in feces are associated with increased adenomagenesis in *in vivo* models. This concomitant increase is also associated with increased expression of *MUC2* and genes involved in responding to DNA damage. Overall, treatment of PIRC rats with a biofilm-competent *Desulfovibrio vulgaris* Hildenborough strain provided for colonization of the host colon regardless of the complex endogenous GM, altered the host GM profile, and modulated adenoma burden.

## Discussion

It has previously been reported that the gut microbiota is associated with differences in disease susceptibility and severity of colon cancer (Cani and Jordan, 2018; Chen and Vitetta, 2018; Ericsson et al., 2015; Ijssennagger et al., 2015; Mendes et al., 2018; Mima et al., 2016; Mima et al., 2015; Redelman-Sidi et al., 2018). Numerous reports provide evidence for the role of bacterial taxa that could be opportunistic pathogens, while otherwise existing as commensals in the colon of patients (Chen et al., 2015; Chung et al., 2018; Dejea et al., 2018; Dejea and Sears, 2016; Johnson et al., 2015; Le et al., 2017; Lee et al., 2014; Sears, 2018). Studies comparing normal epithelial and tumor tissues with culture-independent methods, have shown differences in specific bacterial taxa abundances including *Desulfovibrio* spp (Arthur et al., 2012; Chen et al., 2015; Garrett et al., 2009; Kostic et al., 2013; Marchesi et al., 2011; Mima et al., 2015; Sears and Garrett, 2014; Sekirov et al., 2010; Zackular et al., 2013). These bacteria have been associated with healthy controls in colorectal cancer (CRC) studies, including our own where we saw an increased abundance of these taxa in the group with fewer adenomas (Ericsson et al., 2015; Marchesi et al., 2011; Zackular et al., 2013). Herein, we tested the effect of a *Desulfovibrio* sp. on disease susceptibility via treatment of a preclinical rat model of colon cancer in the context of a complex GM. Unlike other studies where CRC development was studied in germ-free or mono-colonized models we tested the ability of a taxon to colonize a complex developing bacterial community to potentially serve as an improved translatable model for probiotic therapeutics for human disease (Abed et al., 2016; Bullman et al., 2017; Chen et al., 2017; Flynn et al., 2016; Kostic et al., 2013; Sears, 2018; Tlaskalová-Hogenová et al., 2011; Wong et al., 2017; Yang et al., 2018; Zackular et al., 2013).

We found that the biofilm-competent WT *Desulfovibrio vulgaris* Hildenborough strain (De León et al., 2017) and fluorescently labeled JWT733 strain colonized the PIRC rat colons despite the presence of an existing complex GM. Shepherd *et al*. (2018) recently showed that strain engraftment in a complex GM setting could be a function of specific bacterial genes and their corresponding carbohydrate substrate establishing a metabolic niche. In our study, we found that the biofilm-forming DvH strain stably colonized within an endogenous complex community, without the need for altering the carbohydrate composition or the diet, which has been a requirement for other bacteria in other studies (Hildebrandt et al., 2009; Kashyap et al., 2013; Zarrinpar et al., 2014). We detected the presence of JWT733 one-week post treatment via PCR and by fluorescent colonoscopy starting at 2 months of age. DvH colonization was observed with fluorescent JWT733 and WT DvH (without fluorescent marker), suggesting that the presence of dTomato does not impact colonization. DvH colonization was also correlated with a decreased adenoma burden (number and average size), regardless of host sex, when compared to rats treated with buffer or biofilm-deficient DvH.

Biofilms are a critical first-step and required for bacterial colonization in the marine, steel and corrosion industries (Acuña et al., 2006; Dang and Lovell, 2016; Donlan, 2002). The capacity to form biofilms has also been linked to host colonization (Martinez-Gil et al., 2010; Yaron and Römling, 2014). It is likely that the proteins exported by the T1SS for biofilm-formation in the wildtype and the JWT733 strains enabled the bacteria to colonize the PIRC rat colon, thereby creating a protective local environment, i.e. at the mucosa. It is evident from the fecal and mucosal biopsy samples at 2 months of age that the GM profiles among the control, JWT716, and the JWT733 treated groups are significantly different. In the JWT733 rats, the abundance of taxa associated with healthy colonic tissues is suggestive of a mucosal-associated community that is likely protective. Some of these OTUs including *Micrococcus* (Xu and Jiang, 2017), *Bifidobacterium* (Uccello et al., 2012), *Coprococcus* (Shen et al., 2010), *Butyrivibrio* and *Allobaculum* (Tomkovich et al., 2017) have previously been reported to be associated with either healthy stool or tissue samples from control patients in CRC studies. It is noteworthy that these differences were observed between 2 weeks and 2 months, which is likely the time period when adenomas are initiated in the growing intestine.

The GM communities of the fecal samples are significantly different from those observed in the biopsies. We found increased relative abundances of butyrate-producers such as *Faecalibacterium* and *Butyricimonas* in the JWT733 group, correlating with previous studies suggesting that these bacteria prevent tumorigenesis (Guo and Li, 2019; Hibberd et al., 2017; Richard et al., 2018). However, there was also higher relative abundances of bacteria such as *Alistipes* and *Bacteroides* that have been associated with increased tumor burden or with carcinoma samples, indicating that spatial organization in the feces and mucosa may be more important (Feng et al., 2015). At one week of age and at 2 months, both the JWT716 and control groups shared OTUs that were associated with increased CRC and were significantly different from the OTUs found in the JWT733 group. *Roseburia, Lachnospiraceae, Ruminococcaceae* and *Prevotellaceae* have been consistently linked with CRC across many studies (Chen et al., 2012; Dejea et al., 2014; Sun et al., 2017; Zackular et al., 2014). Recently, several groups have demonstrated that the gut mucosal- associated microbiota were differential to those observed in fecal samples (Altomare et al., 2019).

We found JWT733 in both fecal and biopsy samples via quantitative PCR, suggesting either the shedding of growing bacteria from the mucosal associated colonization into the lumen, or the colonization in both the fecal and mucosal environments. The concept that taxa may be populating microhabitats within the GI tract has previously been reported by Donaldson *et al*. (2016). Whereas we observed 40% of the histological sections the of the JWT733-treated colons showed fluorescent bacteria associated with the mucosal layer, the methods of fixation and bacterial FISH are highly technical and likely resulted in an underestimate of the biofilm colonization.

The role of sulfate-reducing bacteria on tumor formation is likely complex. Sulfate-reducing bacteria (SRB), including *Desulfovibrio* spp*, Desulfuromonas*, *Desulfotomaculum*, and *Desulfobacterales* found in the GI tract (Hansen et al., 2011; Leavitt et al., 2016; Stewart et al., 2006; Sun et al., 2017) are known to use sulfates for anaerobic respiration. They release hydrogen sulfide into the lumen (Heidelberg et al., 2004; Landry et al., 2018; Ritz et al., 2017; Yadav et al., 2017). SRB utilize hydrogen (H2) or short-chain fatty acids such as lactate or acetate as electron donors. When sulfate is limited, SRB can form syntrophic communities with methanogens by exchange of H2 or acetate for methanogenesis (Ozuolmez et al., 2015; Timmers et al., 2018). These metabolisms affect the array of metabolites available to the microbiota, including the response of the microbiota to diet (Rey et al., 2013). Various studies have shown that hydrogen sulfide promotes colon cancer cell proliferation (Cai et al., 2010; Hellmich and Szabo, 2015), including energy metabolism and angiogenesis in the colon cancer cell lines (Malagrinò et al., 2019; Szabo et al., 2013). Interestingly, we found that the dissolved sulfide levels were significantly higher in the biofilm-deficient (JWT716) and control groups at two months of age, compared to the JWT733 group despite DvH colonization of the latter. It is plausible that other sources may contribute to this increase in sulfide in the control and JWT716 groups such as the increase of other taxa.

Several mucin-degrading bacteria such as *Akkermansia muciniphila*, *Ruminococcaceae, Ruminiclostridium, Lachnospiraceae, Lachnoclostridium* and *Mucispirillum* were also enriched in the JWT716 and the control rats. In contrast to the report by Croix *et al*. (2011), who found that mucin abundance did not correlate with the abundance of sulfate-reducing bacteria, we found a correlation with higher levels of *MUC2* gene expression. However, they noted that the predominant SRB genus in the gut, *Desulfovibrio,* did not show a concomitant increase with sulfomucins, a phenomenon detected with other taxa. The associated increase of *MUC2* gene expression in the control and JWT716 groups suggests that the degradation of mucin, and its subsequent replenishing led to the release of sulfonated compounds leading to increased H2S production along with other sulfate-reducing bacteria. Additional studies will be required to resolve the species and the correlations observed in our disease model.

Complementary to the principle of higher levels of H2S correlating to higher tumor burden we found higher levels of *HIF1α* and *PTGS2* expression in the control and JWT716 animals. The relative high expression of these genes suggests a hypoxic environment possibly due to the higher level of H2S, a reducing agent (Bianco et al., 2017; Bir et al., 2012; Leschelle et al., 2005; Szabό, 2007, Bocca et al., 2012; Habib et al., 2014; Roberts et al., 2011). Simultaneously, increased expression of *PTGS2* has been associated with GI inflammation and increased susceptibility to colon cancer (Cox et al., 2004; Lobo Prabhu et al., 2014; Ng et al., 2015; Wang and DuBois, 2010; Wiesner et al., 2001). Consequential of the elevated and potentially genotoxic nature of sulfide, we noticed higher levels of expression of DNA damage response genes *MSH2, ATM,* and *MGMT* in the control and JWT716 rats (Jackson and Bartek, 2009; Zhou and Elledge, 2000). This suggests H2S may be causing DNA damage in the proliferating colonocytes. Meanwhile, the exogenous sulfides produced in the JWT733 rats within proximity of the mucosal surface may be protective as shown in *in vitro* and *ex vivo* experiments (Chattopadhyay et al., 2012). Supporting this notion, we found decreased levels of fecal sulfide at 2 months in the rats colonized with DvH compared to that of rats treated with buffer as a control or with biofilm-deficient DvH. From these data, it is not possible to discern changes in sulfide production from changes in sulfide solubility. However, the significance of the spatiotemporal arrangement of the complex GM communities within the lumen and the mucosa may be relevant to understanding the etiology of colon cancer and needs further investigation.

Alternatively, we cannot discount that the type 1 secretion systems (T1SSs) in the biofilm competent DvH strains are exporting a peptide that is responsible for the reduced adenoma burden. The T1SS is necessary for transport of some polypeptides across the bacterial outer membrane. They secrete a wide range of proteins including adhesins, cyclases, metalloprotease-phosphatases, hydrolases, hemolysins etc. (Abby et al., 2016; Delepelaire, 2004; Green and Mecsas, 2016; Morgan et al., 2017). The T1SS ABC transporter in DvH is proposed to export two proteins (designated by their gene loci DVU1012 and DVU1545; De León et al., 2017). Both proteins are annotated as hemolysin-type calcium binding repeat proteins. DVU1012 has a von Willebrand factor A domain which is thought to be involved in cell attachment in eukaryotic cells (De León et al., 2017). These proteins also share similarities with the RTX (repeat-in-toxin) proteins recently reported in *E.coli* as being required for colonization of the urinary tract and kidneys (Vigil et al., 2012). One of the functions of the RTX family of genes is the production of alpha-hemolysin, reported in several Gram-negative bacteria (including *E.coli*) to be capable of causing urinary tract infections and host tissue damage (Bauer and Welch, 1996; Sasaki et al., 2009; Vigil et al., 2012).

Some reports have suggested that hemolysins promote tumorigenesis (Hernández-Luna et al., 2016), while others propose that bacterial hemolysins could be protective against colon cancer (Chowdhury et al., 2011). This disparity in the role of the hemolysins is a potential factor affecting the mechanism of reduced burden in the biofilm-competent strain-treated PIRC rats and warrants further investigation in future studies.

In summary, these data show that the ABC transporter gene in a type 1 secretion system of *Desulfovibrio vulgaris* Hildenborough, required for biofilm formation, is also essential for colonization of the colon in Pirc rats. Whether directly or indirectly, the T1SS is involved in reduced adenoma burden in this model of early onset colon cancer. We show that treatment with a bacterium in a complex GM setting could lead to significant shifts in the community structure and affect host gene expression during this process. This is also the first report demonstrating that reduced dissolved sulfide levels in the feces are associated with reduced adenomagenesis using an *in vivo* model. The high levels of sulfide in feces are associated with higher levels of expression of *MUC2* and DNA damage response genes in a period when adenomas are likely initiated or expanding. Overall, treatment with a biofilm-competent *Desulfovibrio vulgaris* Hildenborough strain altered the host GM profile, where the bacterium colonized the colon regardless of the complex endogenous GM, and ultimately modulated adenoma burden. Our study emphasizes the complex and synergistic interactions, including the possibility that sulfide has different effects contingent on the spatial arrangement of the GM, simultaneously affecting the susceptibility and etiology of colon cancer.

## Methods

### Key resources table

| REAGENT or RESOURCE | SOURCE | IDENTIFIER |
| --- | --- | --- |
| Animal models |  |  |
| F344/NTac-*Apc*<sup>+/am1137</sup>Uwm | Rat Resource and Research Center (RRRC) | RRRC#782<br>RS:0001236 |
| F344/NHsd | Envigo Laboratories | Fischer 344 Inbred Rat<br>RS:0000363 |
| Chemicals, reagents, and other consumables |  |  |
| Ventilated racks | Thoren | Free Standing Housing System |
| Rat cage | Thoren | #4 |
| PicoLab Mouse Diet 20 | LabDiet | #5058 |
| LC medium | Recipe (De León et al., 2017)<br>Components from Fisher Scientific | NA |
| Kanamycin sulfate | Gold Biotechnology | K-120-2 |
| Spectinomycin dihydrochloride pentahydrate | Gold Biotechnology | S-140-25 |
| Vinyl anaerobic chamber | Coy Laboratory Products | Type C |
| MOYLS4 medium | Recipe (De León et al., 2017) | NA |
| G418 sulfate | Gold Biotechnology | G-418-10 |
| L(+)-arginine hydrochloride | Acros Organics | 105000250 |
| Glucose (Dextrose) | Fisher Scientific | D16-500 |
| Glycerol | Fisher Scientific | 56-81-5 |
| Phosphate Buffered Saline, pH 7.3 | Recipe (Ausubel et al., 1994)<br>Components from Fisher Scientific | NA |
| Neubauer counting chamber | Clay-Adams |  |
| Plastic feeding (gavage) needle | INSTECH | FTP-20-38-50 |
| Sterile polyester tipped applicators | Puritan | 25-806-1PD |
| Eppendorf tubes, 1.5 mL | ThermoFisher Scientific | 022 364111 |
| Cysteine | Sigma-Aldrich | C7352 |
| Isoflurane | Vet One | 1000008760 |
| Isothermal pad (Deltaphase) | Braintree | 39DP |
| Brain Heart Infusion Broth, BHIB | Sigma-Aldrich | Cat #53286 |
| Biopsy forceps | Olympus | FB-230U |
| Carbon dioxide | Airgas |  |
| Scalpel blade | Feather | No. 22 |
| Liquid nitrogen | Airgas |  |
| Dulbecco's- Phosphate-Buffered Saline, D-PBS | Sigma-Aldrich | Cat #14190-169 |
| Trypsin | Sigma-Aldrich | Cat #T3924 |
| Collagen | Invitrogen | Cat #A1048301 |
| Methacrylate resin | Electron Microscopy Sciences | Technovit 8100 |
| Ethanol | ThermoFisher Scientific | BP2818 |
| Microscope | Leica | M165FC |
| Petri dish | Celltreat | 229653 |
| Low-melt agarose | IBI Scientific | IB70051 |
| 24-well culture plate | Thermo Scientific | 174931 |
| Paraformaldehyde | Electron Microscopy Sciences<br>Alfa Aesar (16% liquid Ampule) | 15712-S<br>43368 |
| BEEM capsules | Electron Microscopy Sciences | NC9726573 |
| Glucose | Sigma-Aldrich | G8769 |
| Microtome | Dupont Instruments | Sorvall JB-4 |
| Confocal microscope | Leica | SP-8 |
| Serum bottle | Wheaton | 223 738 |
| Glass vial, 2 mL | Fisher Scientific | 03-391-8 |
| Chloroform | Fisher Scientific | C2432 |
| Methanol hyper grade for LC-MS | Merck | Ref 106035 |
| Zinc acetate | MP Bio | 152593 |
| 12 N Hydrochloric acid | Sigma | H1758 |
| N,N-dimethyl-p-phenylenediamine sulfate | Sigma Aldrich | D4139 |
| Technovit® 8100 | Electron Microscopy Systems | Cat # 14654 |
| Carnoy's solution | (Puchtler et al., 1968) | NA |
| FeCl<sub>3</sub>·6H<sub>2</sub>O | Sigma | F2877 |
| Microplate Reader | BioTek | Synergy Mx |
| sodium sulfide nonahydrate | Fisher Scientific | 50-901-13904 |
| Critical Commercial Assays |  |  |
| AllPrep DNA/RNA/Protein Kit | Qiagen | Cat # 80004 |
| NEBNext Ultra Directional RNA Library Prep Kit for Illumina | New England Biolabs | Cat #E7420 |
| PrimeTime® Predesigned qPCR probe | Integrated DNA Technologies | Cat.No. Hs.PT.58.46475178.g |
| SYBR™ Green qPCR SuperMix | Sigma-Aldrich | KCQS02-1250RXN |
| iTaq™ Universal SYBR® Green Supermix | Bio-Rad | 1725131 |
| Experion RNA StdSens analysis kit | Bio-Rad | 7007104 |
| Sso Advanced Universal SYBR Green Supermix | Bio-Rad | 172-5131 |
| SuperScript™ III First-Strand Synthesis System | Invitrogen | 18080051 |
| Qubit dsDNA HS Assay Kit | ThermoFisher Scientific | Q32854 |
| Deposited data |  |  |
| 16S rRNA gene sequencing | This paper | PRJNA495020 |
| Experimental models: Organism/Strains |  |  |
| Bacterial strains and plasmids | This paper (Supplementary Table 1) | NA |
| Oligonucleotides |  |  |
| Primers and probes for qPCR and bacterial strain generation | This paper (Supplementary Table 2) | NA |
| Software and Algorithms |  |  |
| Qiime | (Caporaso et al., 2010) | 23ersion 1.9 |
| SILVA database | (Quast et al., 2013) | version 132; December 13, 2017 |
| Leica Application Suite | Leica | 23ersion 4.2 |
| GraphPad Prism 7 | GraphPad | version 7 for Windows |
| DESeq2 1.16.1 | (Callahan et al., 2016) | http://www.bioconductor.org/packages/release/bioc/html/DESeq2.html |
| Correlogram | (Wei and Simko, 2017) | *corrplot* package Version 0.84 |
| PAST | (Hammer et al., 2001) | NA |
| R | https://www.r-project.org/ | Version 3.1.4 |
| Bowtie 2 | (Langmead and Salzberg, 2012) | Version 2.2.9 |
| Other |  |  |
| Syringe 5ml | BD Biosciences | 309646 |
| Aluminium Crimp | Fisher Scientific | 12373680 |
| Needle 25 gauge (length: 5/8") | BD Biosciences | 305122 |
| 1ml TB syringe w/ 26 gauge (length: 3/8") | BD Biosciences | 309625 |
| Syringe 3ml lure lock | BD Biosciences | 309657 |
| Septum penetration needles (deflected point; 18 gauge; 6") | Cadence Inc. | 4195 |
| Sterile swabs | ThermoFisher Scientific | 165KS100 |
| Cryovials | ThermoFisher Scientific | 366656PK |
| Bioanalyzer | Agilent | 2100 |
| Realtime PCR Instrument | Bio-Rad | CFX-384 |

### LEAD CONTACT AND MATERIALS AVAILABILITY

Further information and requests for resources and reagents should be directed to and will be fulfilled by the Lead Contact, James Amos-Landgraf.

### EXPERIMENTAL MODEL AND SUBJECT DETAILS

#### Animal husbandry and housing

PIRC rats were generated by crossing male F344/NTac-*Apc^+/am1137^*Uwm (Rat Resource and Research Center #782) rats with wild-type female F344/NHsd rats obtained commercially from Envigo Laboratories (Indianapolis, IN). Animals were housed in micro-isolator cages on ventilated racks (Thoren, Hazleton, PA) and acclimated for a week in groups prior to breeding. Cages were furnished with paper chip bedding and were fed irradiated 5058 PicoLab Mouse Diet 20 (LabDiet, St. Louis, MO). Rats had *ad libitum* access to water purified by sulfuric acid (pH 2.5-2.8) treatment followed by autoclaving. Fecal samples were collected with aseptic methods to be used as a reference from all breeders prior to cohousing. After allowing for one day of mating to establish timed pregnancies, females were moved to new cages and individually housed thereafter.

#### Ethics Statement

The study reported here was conducted in accordance with the guidelines established by the Guide for the Use and Care of Laboratory Animals and the Publish Health Service Policy on Human Care and Use of Laboratory Animals. All studies and protocols (#6732 and #8732) were approved by the University of Missouri Institutional Animal Care and Use Committee.

#### Detailed methods

##### Genotyping and animal identification

Pups were ear-punched prior to weaning at 13 days of age. DNA was extracted with the “HotSHOT” genomic DNA preparation method and was used for a high resolution melt (HRM) analysis to differentiate wildtype rats from those carrying the *APC* mutation (PIRC) as described previously (Ericsson et al., 2015; Truett et al., 2000).

##### Bacterial strains, media, and growth conditions

All strains and plasmids used in this study are presented in Supplementary Table 1 and are available upon request. Methods for growth of *Escherichia coli* and DvH cultures and for plasmid generation in *E. coli* were performed as described previously (De León et al., 2017). Briefly, *E. coli* cultures were grown at 37 °C on LC medium containing either kanamycin or spectinomycin (50 or 100 μg/ml, respectively; Gold Biotechnology, Inc., St. Louis, MO) and used as the host for recombinant plasmids made via sequence and ligation-independent cloning (SLIC; Li and Elledge, 2007). The primers used to amplify fragments for the SLIC reaction and to confirm the plasmid via sequencing are shown in Supplementary Table 2. DvH cultures were grown at 30 °C in an anaerobic growth chamber (approximately 95 % N_2_ and 5 % H_2_; Coy Laboratory Products, Inc., Grass Lake, MI) in liquid or solidified lactate/sulfate medium supplemented with 1 % (wt./vol) yeast extract (MOYLS4; De León et al., 2017). Where indicated, G418 (400 μg/ml; Gold Biotechnology, Inc), spectinomycin (100 μg/ml), or L(+)-arginine hydrochloride (126.5 μg/ml; Acros Organics, New Jersey; per Davis RW, 1990) were added to the DvH cultures. DvH cultures were routinely inoculated onto LC plates containing 40 mM glucose and incubated aerobically at 30 °C for at least two days to ensure there was no aerobic contamination.

Wild-type DvH was received from American Type Culture Collection (ATCC) in ca. 2007 and was maintained at Montana State University before being sent to the University of Missouri in 2013. This strain was specified as DvH-MT previously for clarification (De León et al., 2017), but is considered to be the wild-type strain and is called DvH here. DvH and DvH-MO contain 30 deviations from the original genome sequencing (Heidelberg et al., 2004) [NCBI GenBank accession no. NC_002937.3] that are likely errors in the original sequencing; 29 of these were reported previously (De León et al., 2017) [NCBI BioProject PRJNA392176] and one additional difference has since been identified [C(10) to C(11) at genome location 3,254,681]. Our wild-type DvH culture has 17 SNPs detectable in the population but none that are in >50% of the population (range 15-45%). DvH-MO is a spontaneously biofilm-deficient strain that contains an additional 12 variants unique to this culture and at or near 100% frequency within the population. One of these 12 variants, a single nucleotide change in the ABC transporter of the type I secretion system (DVU1017) is the cause of biofilm deficiency in this strain (De León et al., 2017). The culture of DvH-MO used in this study was made by combining three isolated colonies after the culture underwent single colony isolation to remove possible rare variants, including genetic revertants, in the population. JWT716 is a markerless-deletion mutant of DVU1017 derived from wild-type DvH and has been described previously (De León et al., 2017).

In preparing cultures to be introduced into the rat gastrointestinal tract, one ml of a frozen stock stored at -80°C in 10% (vol/vol) glycerol in growth medium was thawed, inoculated into 10 ml of MOYLS4 medium, and incubated anoxically at 30 °C. After approximately 24 h, the culture reached an optical density of 0.8 at 600 nm (late logarithmic phase). The cells were pelleted by centrifugation at 3696 x g for 12 min and the pellet was washed with 10 ml of sterile phosphate buffered saline (PBS) pH 7.3 (Ausubel et al., 1994). Centrifugation was repeated and the pellet was resuspended in approximately 10 ml of PBS to yield a final cell concentration of approximately 5 x 10^8^ cells/ml which was confirmed by direct cell count in a Neubauer counting chamber (Clay-Adams Co. New York).

#### Fluorescent strain (JWT733) construction

To generate a fluorescent DvH lacking antibiotic resistance markers, arginine prototrophy was used as a selectable phenotype. *argH* (encoding arginosuccinate lyase; DVU1094) is the last gene of an operon encoding three genes putatively involved in arginine biosynthesis. A plasmid, pMO7722, was constructed containing a gene encoding neomycin phosphotransferase II with its native promotor and conferring kanamycin resistance. To create a marker exchange deletion of 695 bp at the 3′ end of *argH* (full length is 1383 bp), a sequence internal to *argH* (165-688 bp) was placed upstream of the antibiotic resistance cassette and a 511 bp sequence from downstream of *argH* was placed downstream of the cassette. This plasmid, pMO7722, was transformed into wild-type DvH via electroporation as described previously (Keller et al., 2009). Selection of the marker-exchange deletion mutant in which the 3′ end of *argH* (689-1383bp) was replaced with the kanamycin resistance cassette and was auxotrophic for arginine was accomplished by selection in solidified MOYLS4 containing G418 and arginine. Resistance to the kanamycin analog G418, sensitivity to spectinomycin, and arginine auxotrophy were confirmed by growth studies and the genome structure was confirmed by Southern blot. One isolate was obtained and designated JWT726 to be used for the introduction of gene(s) of choice by prototrophic selection. Subsequently, to introduce a fluorescent marker into JWT726 (by the same transformation methods), pMO7743 was constructed to reintroduce the 3′ end of *argH* along with the fluorescent marker, *dTomato* (Shaner et al., 2004). After electroporation, the cells recovered at 30 °C in one ml of MOLS4 (MOYLS4 without yeast extract) for 24 h and were then diluted 10-fold with MOLS4 to select for cells capable of synthesizing arginine. After four days, growth was observed and serial dilutions of this culture were embedded into solidified MOYLS4 for single colony isolation. Colonies showing fluorescence under the microscope were selected for phenotypic confirmation of G418 and spectinomycin sensitivity as well as arginine prototrophy. Upon genomic structure confirmation by Southern blot, one isolate was designated JWT733.

#### Bacterial treatment and necropsy scheme

F344-*Apc^+/am1137^* PIRC rats were used for all the experiments (Fig.1B). For the preliminary study, on days 14 and 15 of age, male and female PIRC rats were treated with 200 µL of ∼10^8^ cells per ml of either wild type DvH or DvH-MO suspended in anaerobic PBS (phosphate buffered saline) via oral gavage. All rats were subsequently weaned from the mothers at 21 days of age. Adenoma growth was confirmed through colonoscopies every month starting at two months of age (Irving et al., 2014b). At 4 months of age, animals were sacrificed post-disease onset as described previously (Ericsson et al., 2015). For the follow-up study to determine if the point mutation in the wild-type DvH strain influenced the adenoma phenotype, we followed the same protocol and timeline as the above. However, rats were treated with either JWT733 or JWT716 strains of DvH. Rats from the control group were simultaneously gavaged with 200 µL of anaerobic PBS (pH 7.3) to serve as a negative and gavage control.

#### Fecal Collection

Sterile swabs (ThermoFisher Scientific, Waltham, MA) were used to obtain a pre-treatment fecal sample on day 13 of age. Fecal samples from adult rats at weaning (21 days of age) and post-weaning (starting at day 30 of age) were collected by placing the animal in a clean, sterile cage without bedding. Fecal samples were collected at 1-week post treatment and monthly starting at 1 month of age. Freshly evacuated feces were speared with sterile toothpick or forceps and placed into a sterile Eppendorf tube. All samples were collected into cryovials (ThermoFisher Scientific) and stored at -80 °C until processing for 16S rRNA gene analysis.

#### Fecal DNA extraction, 16S library preparation and sequencing

Fecal samples were pared down to 65 mg with a sterile blade and then DNA was extracted by the method described previously (Ericsson et al., 2015). Amplification of the V4 hypervariable region of the 16S rRNA was performed at the University of Missouri DNA core facility (Columbia, MO) also, as previously described (Ericsson et al., 2015). Briefly, bacterial genomic DNA was used for amplification of the V4 hypervariable region with universal primers (U515F/806R) flanked by Illumina standard adapter sequences and the products were pooled for sequencing on the Illumina MiSeq platform. Samples with more than 10,000 reads were used for assembly, binning and annotation with QIIME v1.9 including trimming and chimera removal as described previously (Montonye et al., 2018). The data obtained from the pre-treatment samples, i.e. swabs, did not meet the criteria for the listed sample inclusion and so were not included in the analyses. Based on 97% nucleotide identity, contigs were assigned to operational taxonomic units (OTUs) via *de novo* OTU clustering. These OTUs were annotated with BLAST (Altschul et al., 1990) against the SILVA database 132, released on December 13, 2017 (Hart et al., 2015; Quast et al., 2013).

#### Colonoscopy

Rats were anaesthetized with isoflurane (3 % vol/vol) and placed on a heating pad to maintain body temperature. Minimal use of sterile PBS (∼1ml) was used to clear colonic contents helping to lubricate and remove any fecal material. Endoscopic video and images were recorded as previously described (Irving et al., 2014b). Colonic tissue samples from normal epithelium (3 mm^3^), i.e. non-tumor tissues adjacent to tumors, were collected at two months of age, with biopsy forceps (FB-230U, Olympus, USA).

#### Necropsy, normal epithelium and tumor tissue collection

All animals were humanely euthanized with CO_2_ administration and necropsied at sacrifice. The small intestine and colon from the rats were placed onto absorbent paper and then opened longitudinally. With a sterile scalpel blade (Feather, Tokyo, Japan), normal colonic epithelium tissues were scraped from the top, middle and distal regions of the colon. Tumors in the same locations were collected by resecting half of the tumor. All tissues were flash-frozen in liquid nitrogen and stored at -80 °C. Remaining intestinal tissues were then fixed overnight in Carnoy solution (Puchtler et al., 1968),which was replaced with 70% (vol/vol) ethanol for long term storage until adenoma counting was performed.

#### Methacrylate embedding, sectioning and confocal microscopy

The following protocol was modified from Mark Welch *et al* (2017). Excised tissues, described above were gently coated with 0.5% (wt./vol) low melting point agarose (ThermoFisher Scientific), placed into a well in a 24-well cell culture plate (ThermoFisher Scientific). The tissues in agarose were allowed to harden for 2 hours at 4 °C. The samples were then removed from the agarose and fixed in 2% (vol/vol) paraformaldehyde for 12 hours at 4 °C. Samples were washed with PBS, and again coated with 0.5% (wt./vol) molten agarose. Excess agarose was trimmed before embedding into methacrylate resin with the Technovit^®^ 8100 system (Electron Microscopy Systems, Hatfield, PA) per manufacturers’ guidelines with the following modifications: samples were dehydrated with acetone for one hour at 4 °C, with repeated changes of acetone, until the solution remained clear; the sample was then covered with the infiltration solution to set overnight at 4 °C. Polymerization was done by adding 400 µL of embedding solution to the samples within BEEM capsules (Electron Microscopy Services) and allowed to set overnight at 25 °C in an anaerobic chamber due to the oxygen sensitivity of the embedding solution. The samples were sectioned to 5 µm thickness with a Sorvall JB-4 Microtome (Dupont Instruments, Connecticut, USA). We subsequently visualized via confocal microscopy with an SP-8 system (Leica Microsystems, Buffalo Grove, IL). Fluorescent in situ hybridization (FISH) was performed as described by Mark Welch *et al* (2017) with probes listed in Supplementary Table 2.

#### Sulfide assay

Three fecal pellets from each rat were collected immediately after evacuation and each was placed with sterile forceps into a 10 mL sterile serum bottle under anaerobic conditions (Fisher Scientific, Pittsburgh, PA) as technical triplicates. Each serum bottle contained a smaller 2 mL vial (Fisher Scientific) with 1mL of freshly prepared 2% (wt./vol) zinc acetate. Prior to adding fecal samples, the serum bottles and vials were equilibrated with the atmosphere of the anaerobic chamber to remove oxygen by leaving them for 48 hours in an anaerobic chamber. After introduction of fecal pellets, the bottles were flushed again with nitrogen to maintain an anaerobic environment. Thereafter, the vials and bottles were handled inside the chamber, and sealed to maintain a non-oxygenated environment within the bottles. Cline’s sulfide assay (Cline, 1969) was modified to determine the levels of sulfide dissolved in fecal samples spectrophotometrically at 670 nm by a passive capture technique modified from Ulrich *et al* (Ulrich et al., 1997). Briefly, 0.3 mL of 12 N hydrochloric acid was used to drive dissolved sulfides from fecal material into gaseous form to be captured passively by the zinc acetate solution. Following 24 hours of passive capture of the sulfides, the zinc acetate solution containing captured sulfide was removed from the vial and 200 µL was treated with 16 µL of Cline’s reagent within a white, 96-well, optical bottom plate (ThermoFisher Scientific). Cline’s reagent was comprised of 8g of *N,N*-dimethyl-*p*- phenylenediamine sulfate and 12g of FeCl_3_·6H_2_O in 500 mL of 50% (vol/vol) hydrochloric acid. Following a 20 min incubation at room temperature, the absorbance at 670 nm was measured in a BioTek Synergy Mx microplate reader (BioTek Instruments Inc., Winooski, VT). A calibration curve of standards was established with sodium sulfide nonahydrate (Na_2_S·9H_2_O) in 2% (wt./vol) zinc acetate to determine the concentration of sulfide per sample. The sulfide was normalized to the weight of each fecal pellet.

#### RT-qPCR and gene expression analysis

Total RNA was extracted from biopsies of normal colonic tissues with the Allprep DNA/RNA/Protein Mini kit (Qiagen, Germantown, MD) and reverse-transcribed into cDNA with the SuperScript III First-Strand Synthesis System (Invitrogen, Carlsbad, CA) with the standard described protocol for each kit. Prior to cDNA conversion, the quality of the RNA was assessed by the Experion RNA StdSens analysis kit (Bio-Rad, Hercules, CA). All samples below the RNA

Quality Indicator (RQI) of 7 were excluded from gene expression experiments and analysis. The cDNA for all samples were made from the same concentration of mRNA, i.e. we normalized the input mRNA concentration to that of the lowest mRNA concentration from all samples. The data were additionally normalized to a house-keeping gene (GAPDH) in the subsequent analysis. Reverse-transcriptase quantitative polymerase chain reaction (RT-qPCR) for mRNA expression was used to assay the following host genes: *MSH2* (MutS Homolog2)*, ATM* (Ataxia Telangiectasia Mutated Serine/Threonine kinase)*, MGMT* (O-6-Methylguanine-DNA Methyltransferase)*, HIF1α* (Hypoxia Inducible Factor 1 Subunit Alpha)*, NOX4* (NADPH Oxidase 4)*, PTGS2* (Prostaglandin- Endoperoxide Synthase 2), and *CAR1* (Carbonic Anhydrase 1)*. GAPDH* (encoding glyceraldehyde phosphate dehydrogenase) was used as the housekeeping gene for host gene expression (Barber et al., 2005; Li et al., 2017), while the 16S rRNA gene and *gyrB* (encoding gyraseB) were used as bacterial housekeeping genes (Rocha et al., 2015). *MUC2* (Mucin 2) expression was determined with a PrimeTime® Predesigned qPCR probe (Cat.No. Hs.PT.58.46475178.g, Integrated DNA Technologies, Coralville, IA). *GAPDH* was used as the housekeeping gene for the *MUC2* assay. RT-qPCR was set up with a SYBR^TM^ Green qPCR SuperMix (Thermo Fisher Scientific) in quadruplicate reactions per primer or probe set, per sample. The final PCR mixture contained one µL each of forward and reverse primers (final concentration of 100 nM), 5 µL of 2X SYBR^TM^ Green qPCR SuperMix, 2 µL of sterile water and 1 µL of cDNA at 40 ng/µL. For the *MUC2* assay, the SYBR supermix was replaced with iTaq™ Universal SYBR® Green Supermix (Life Technologies, Carlsbad, CA). The reaction protocol was carried out with an initial incubation of 10 min at 95 °C followed by 40 cycles of the following: denaturing at 95 °C for 15 s, annealing and elongation at 60 °C for 1 min. The forward and reverse primers used are listed in Supplementary Table 2.

#### Tumor counts and size measurements

At the terminal time point of 4 months of age, 0.5 cm sections of the colon were resected as a cylinder prior to splaying open and embedded with a methacrylate resin (Technovit 8100, Electron Microscopy Sciences, Hatfield, PA). The remaining colon sections were cut longitudinally and fixed on absorbent paper with Carnoy solution. Tumor multiplicity was determined by a double-blind gross counting of colonic tumors visualized with a Leica M165FC microscope (Leica, Buffalo Grove, IL) at 5X magnification (Amos-Landgraf et al., 2014; Ericsson et al., 2015; Irving et al., 2014a). Briefly, the small intestine and colonic tissues were laid flat in a 100 mm x 15 mm petri dish (Sycamore Life Sciences, Houston, TX) and covered with 70% (vol/vol) ethanol (ThermoFisher Scientific, Waltham, MA) to prevent tissue drying and visually counted. Tumor sizes were measured with the Leica Application Suite 4.2, after capturing post-fixed images as previously described (Ericsson et al., 2015).

#### Quantification and statistical analysis

All statistical details of the experiments are described in the figure legends and the within the description of the respective paragraph. Additional descriptions of software, packages and algorithms employed for the analysis of 16S rRNA sequencing data, and correlation analyses are described below:

#### Statistical analyses and figures

All statistical analyses and graphs (except Fig.1) were prepared through GraphPad Prism version 7 for Windows (GraphPad Software, La Jolla, CA). *p*-values were considered significant for values less than 0.05 unless otherwise indicated. Analysis of Variance (ANOVA) with a Tukey’s post-hoc test was used to identify differential groups. For OTUs comparison and significance testing DESeq2 was used as described by Callahan *et al*. (2016). Correlations were performed with the linear regression module available through GraphPad Prism v7. Correlation of tumor counts with OTUs depicted as a correlogram were performed in the *corrplot* package v.0.84 (Wei and Simko, 2017) of R software v.3.1.4, with a Pearson correlation coefficient. Heatmaps and PCoAs (principal component analyses) were generated with the open source PAST (Paleontological Statistics) 3.14 software (Hammer et al., 2001).

#### Data and code availability Availability of Data and Material

The raw 16S rRNA gene sequencing files generated for the study, and from which the data were analyzed are available at the associated NCBI BioProject number PRJNA495020. The code used for the analyses is provided as a Supplementary material

## Author contributions

Experiments were conceived and designed by Susheel Bhanu Busi, Kara B. De León and James Amos-Landgraf. Kara B. De León and Judy Wall designed and developed the mutant strain of *Desulfovibrio vulgaris* Hildenborough and contributed reagents including providing bacterial cultures at time of inoculation. Daniel Montonye gavaged the rats with the DvH strains. Data were analyzed by Susheel Bhanu Busi. All authors contributed to writing the paper.

## Acknowledgements

The authors wish to thank Dr. Pamela J.B. Brown and Jeremy J. Daniel at the University of Missouri for kindly providing pSRKKm-tdTomato; Grant M. Zane for the idea of using prototrophy as a selection when introducing genes into genomes; Nathan Bivens and the MU DNA Core for assistance with 16S rRNA sequencing; Bill Spollen, Christopher Bottoms and the MU Informatics Research Core Facility for assistance with software installation for data analysis; Dr. Aaron Ericsson and Dr. Craig Franklin at the MU Metagenomics Center; and the Rat Resource and Research Center and MU Office of Animal Resources and their staff for assistance with animal husbandry.

## Funding

This research was funded by grants from the University of Missouri to Dr. James Amos-Landgraf (Startup-funding) and the MU College of Veterinary Medicine COR grant awarded to Dr. Amos- Landgraf (2017).

## Conflict of interest statement

The authors declare that the research was conducted in the absence of any commercial or financial relationship that could be construed as a potential conflict of interest.

---

**Supplementary figure 1.**
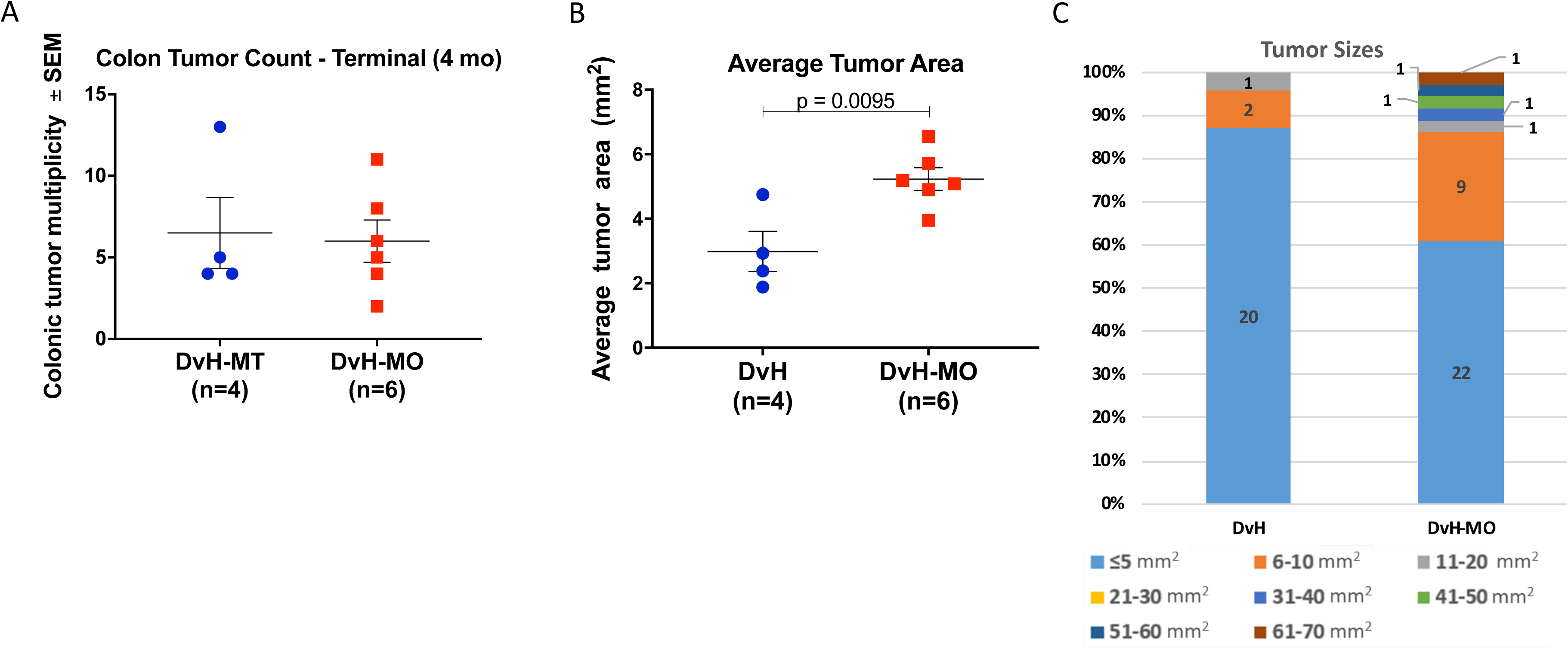
Treatment of PIRC rats with biofilm -competent and –deficient *Desulfovibrio vulgaris* Hildenborough. (A) Colonic tumor count and (B) average tumor size at 4 months. *p*-values were calculated via a Student’s *t-*test and those below 0.05 were considered to be significantly different between groups. (C) Bar graph of the differential tumor sizes observed in the wild type DvH and DvH-MO treated PIRC rats at 4 months. Error bars in all figures indicate standard error of the mean (±SEM). Individual tumor sizes from every rat were plotted in (C), whereas the average tumor count shown in figure (A)

**Supplementary figure 2.**
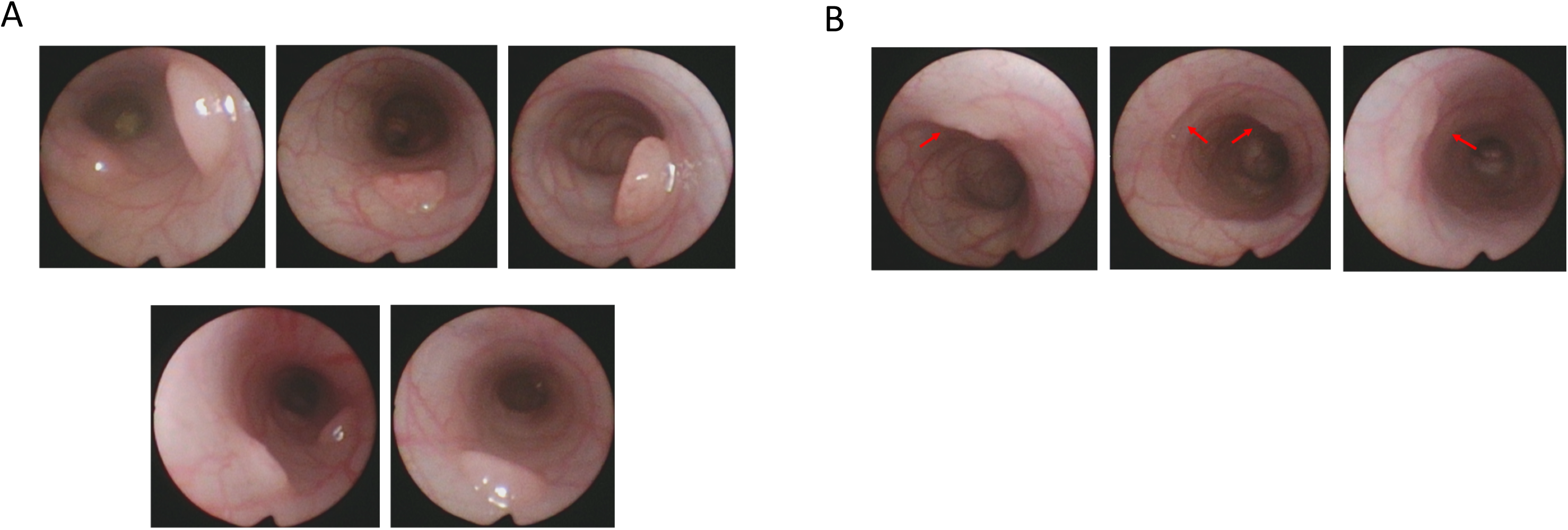
Adenoma images via colonoscopy in DvH-treated PIRC rats. (A) Representative images of adenomas in DvH-MO treated rats indicating larger tumor sizes acquired at 4 months of age (sacrifice). Images were obtained from five different animals. (B) Images are representative of the small lesions observed in the wild type DvH group, obtained from three different animals.

**Supplementary figure 3.**
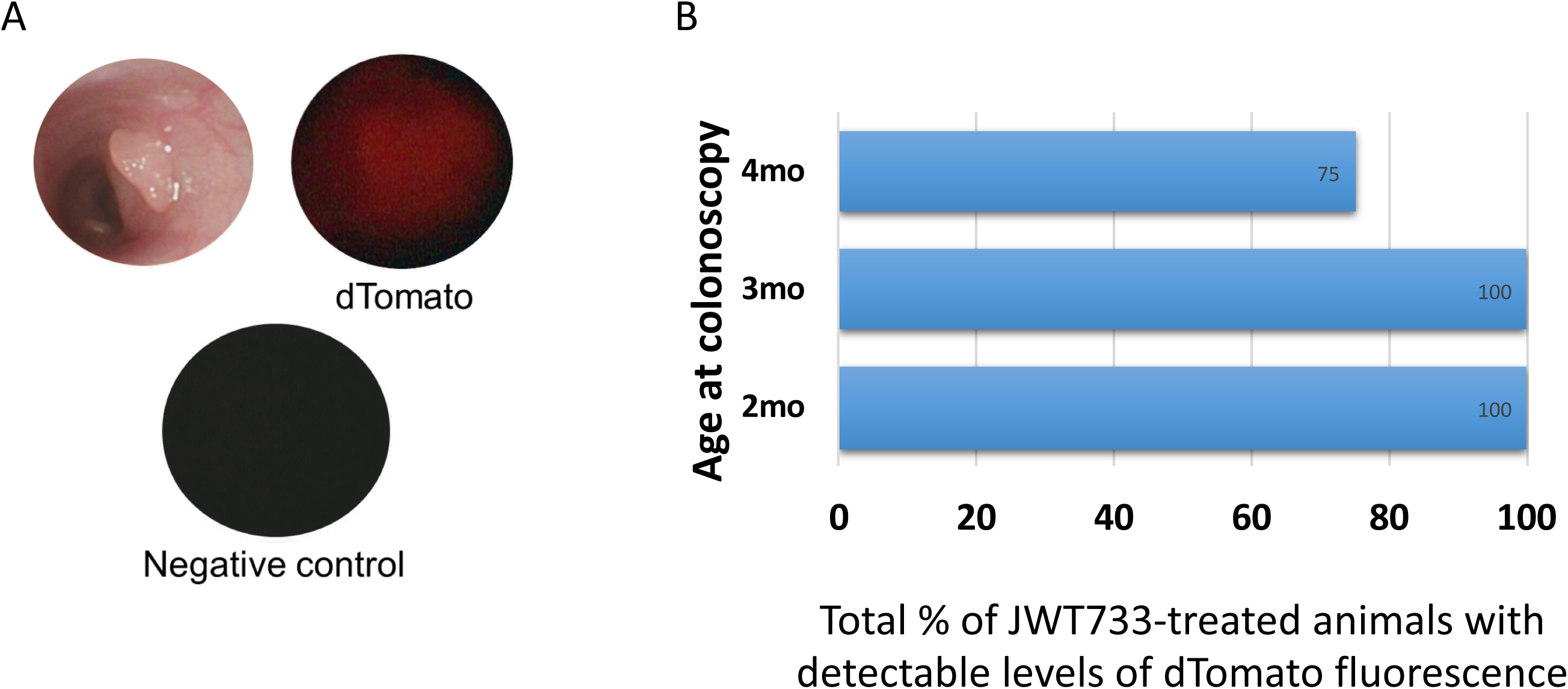
Colonoscopy of fluorescent, biofilm-competent strain-treated rats. (A) Representative images of colonoscopy with white light, dTomato fluorescence and negative controls to determine percent detection of fluorescent in all rats treated with JWT733. (B) Barplot indicates the total percentage of animals treated with biofilm-forming DvH strain, with detectable levels of dTomato fluorescence.

**Supplementary figure 4.**
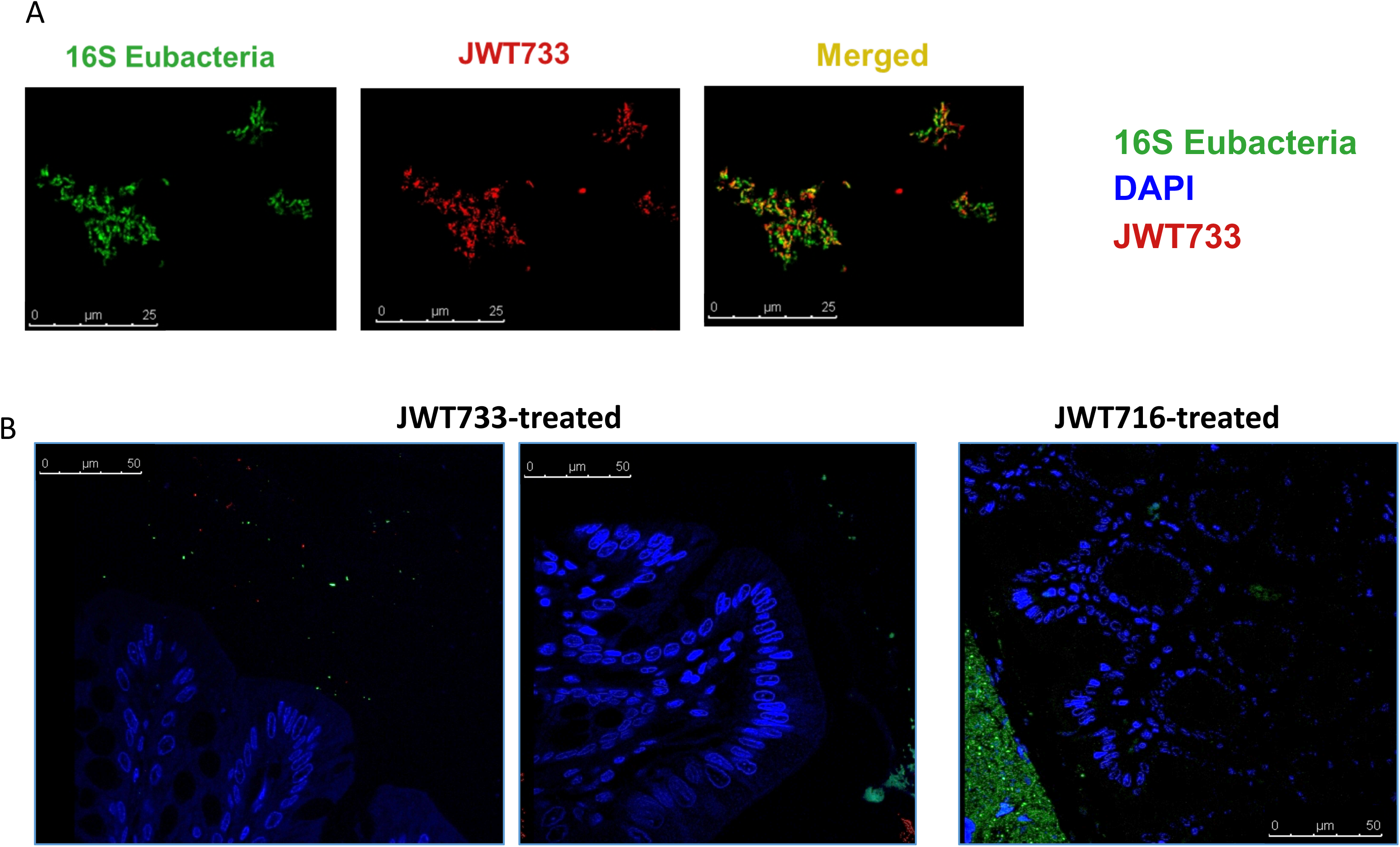
Fluorescent in-situ hybridization (FISH) and confocal microscopy assessing biofilm formation in vivo in the JWT733 treated rats. (A) Confocal microscopy images to detect fluorescent, biofilm-competent JWT733 strain. Representative images of positive controls for 16S Eubacteria and JWT733. (B) Representative images of the JWT733- and JWT716- treated colonic segments assessed for presence of bacteria. JWT733, n=13 and JWT716, n=12.

**Supplementary figure 5.**
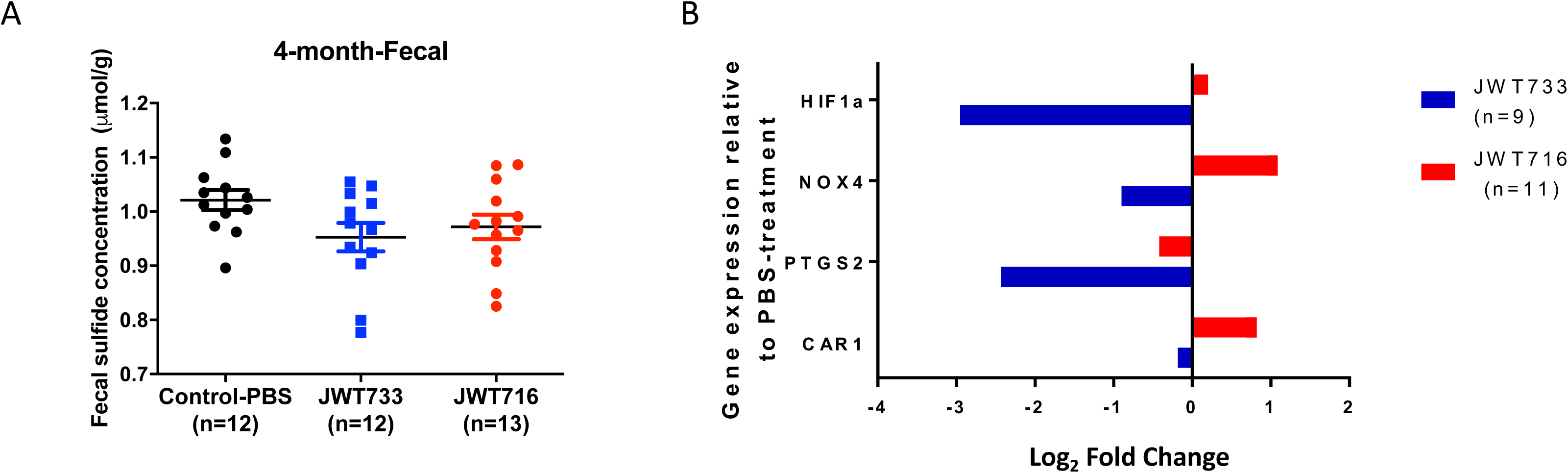
Sulfide assay, RT-qPCR analysis of gene expression in Control, JWT733 and JWT716 groups. (A) Fecal sulfide concentration measured by Cline assay at 4 months of age in the control and treatment groups. p-values below 0.05 were considered to be significantly different between groups. (B) Relative gene expression measured using RT-qPCR with respect to the control (anaerobic-PBS) group determined for inflammation and hypoxia-related genes in all three groups, i.e. Controls (n=8), JWT733 (n=9) and JWT716 groups (n=11). All expression is normalized to *GAPDH* and subsequently to that of the control animals.

**Supplementary figure 6.**
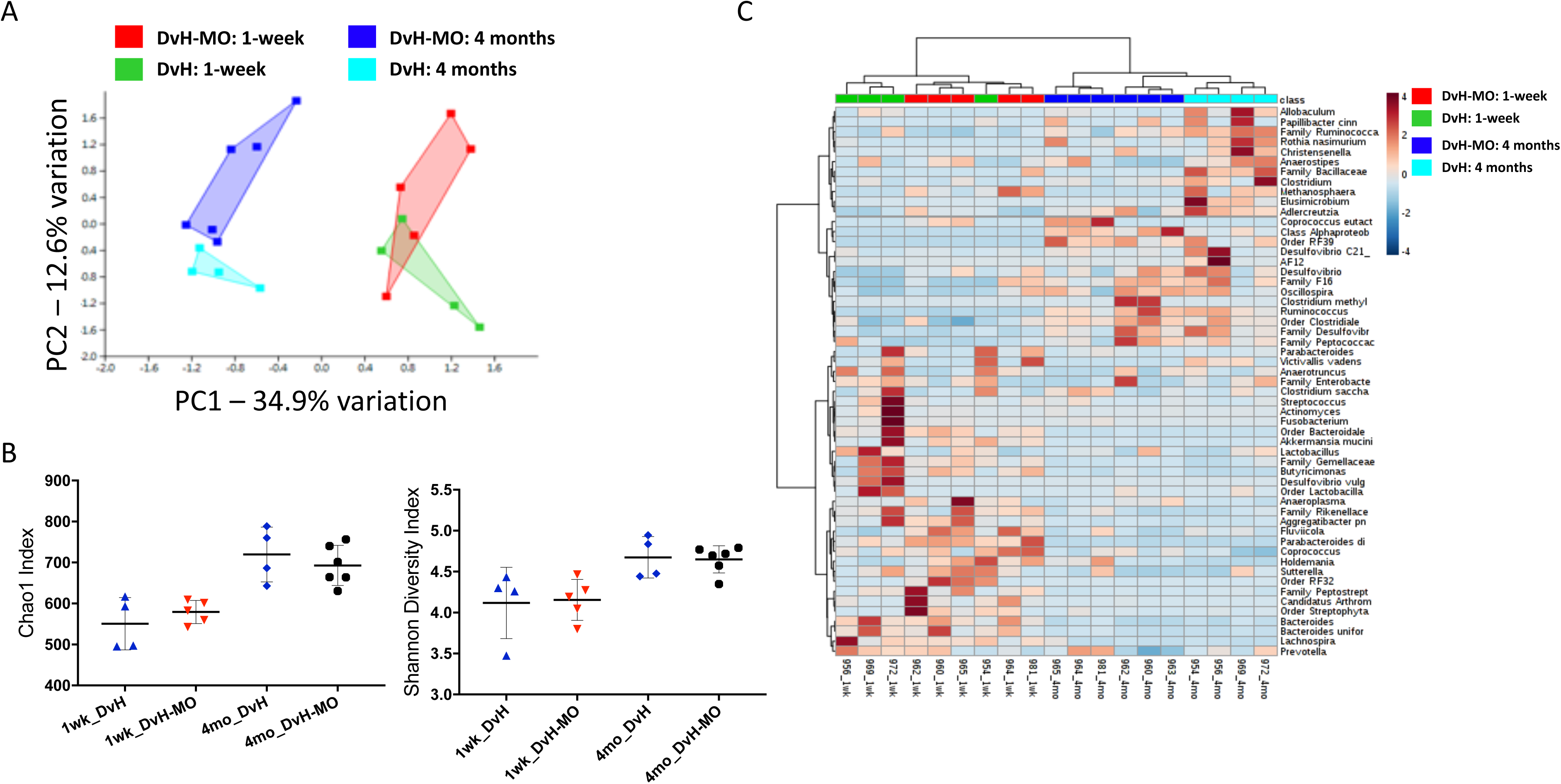
16S rRNA gene sequencing analysis of DvH treatment. (A) Principal component analysis (PCA) indicating the differential complex GM profiles observed in the wild-type DvH (n=4) and DvH-MO (n=6) groups at 1 week (green: wild-type, red: DvH- MO) and 4 months (light blue: wild-type, dark blue: DvH-MO) of age. PERMANOVA (*F*=4.45, *p*=0.0001) was used to determine significance differences in GM profiles. A *p*-value less than 0.05 was considered to be significant. Post-hoc analysis is listed under Supplementary Table 3. (B) Richness (Chao1) and diversity (Shannon) indices were measured for the same time points. (C) Heatmap analysis using Euclidean distances coupled with Ward’s algorithm was performed, identifying the top 55 OTUs (operational taxonomic units). Error bars in all figures indicate standard error of the mean (±SEM).

**Supplementary figure 7.**
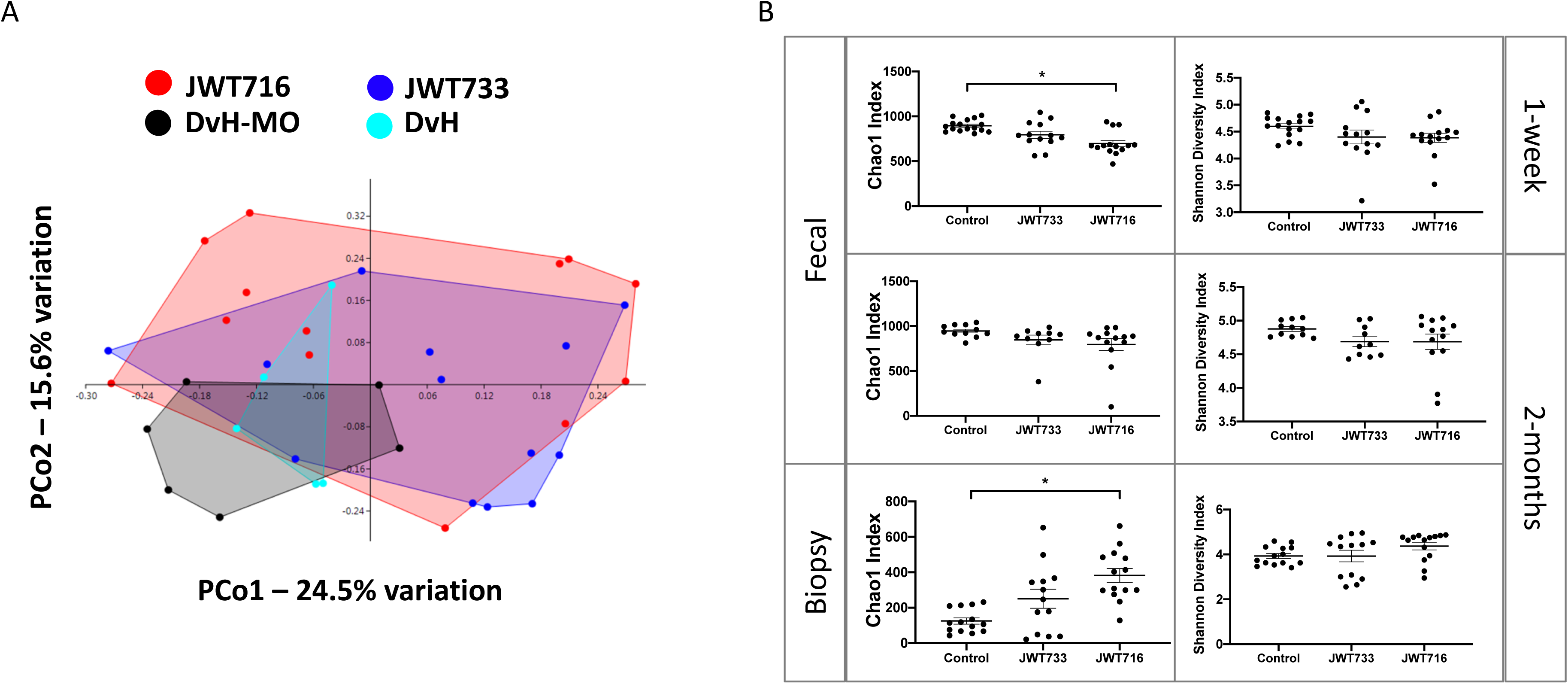
16S rRNA gene sequencing analysis of fecal and biopsy samples. (A) Principal Component Analysis (PCoA) plot depicting the fecal 16S rRNA gene sequencing dissimilarities between the DvH-treated groups based on the Bray-Curtis distance matrix. Post- hoc analysis indicating the differences between individual groups is listed under Supplementary Table 4. Each symbol represents the GM community from the fecal sample of a single rat at 2 months of age. (B) Richness and diversity indices, Chao1 and Shannon respectively, were determined for both fecal and biopsy samples at 1-week after treatment, and 2 months of age. Dot plots represent individual values for each rat. Significance was determined using a Two-way ANOVA, with Student-Neuman Keul’s post-hoc analyses. *represents *p*-value less than 0.05.

**Supplementary figure 8.**
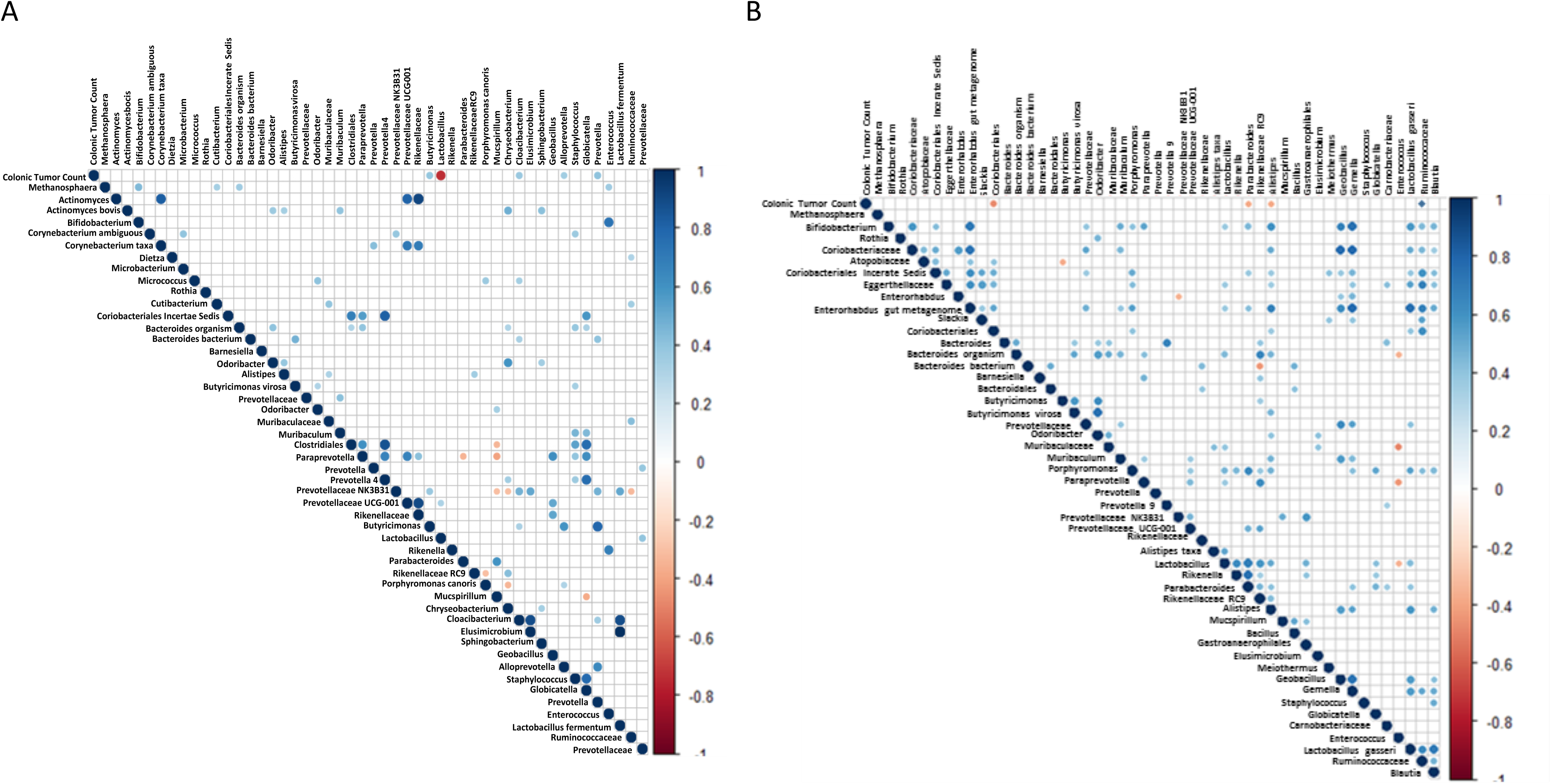
Correlogram analyses of OTUs at one-week and 2-months post treatment vs tumor number at 4 months of age, respectively. (A) Correlogram showing the correlations (Pearson’s, *p*<0.05) between OTUs at one-week post- treatment with colonic tumor counts determined after sacrifice at 4 months of age. Color of the dot indicates positive (blue) or negative (red) correlation. Size of the dot represents the mean relative abundance of each OTU. (B) Correlogram of OTUs from 2-month fecal samples and 4-month colonic tumor multiplicity is depicted.

**Supplementary figure 9.**
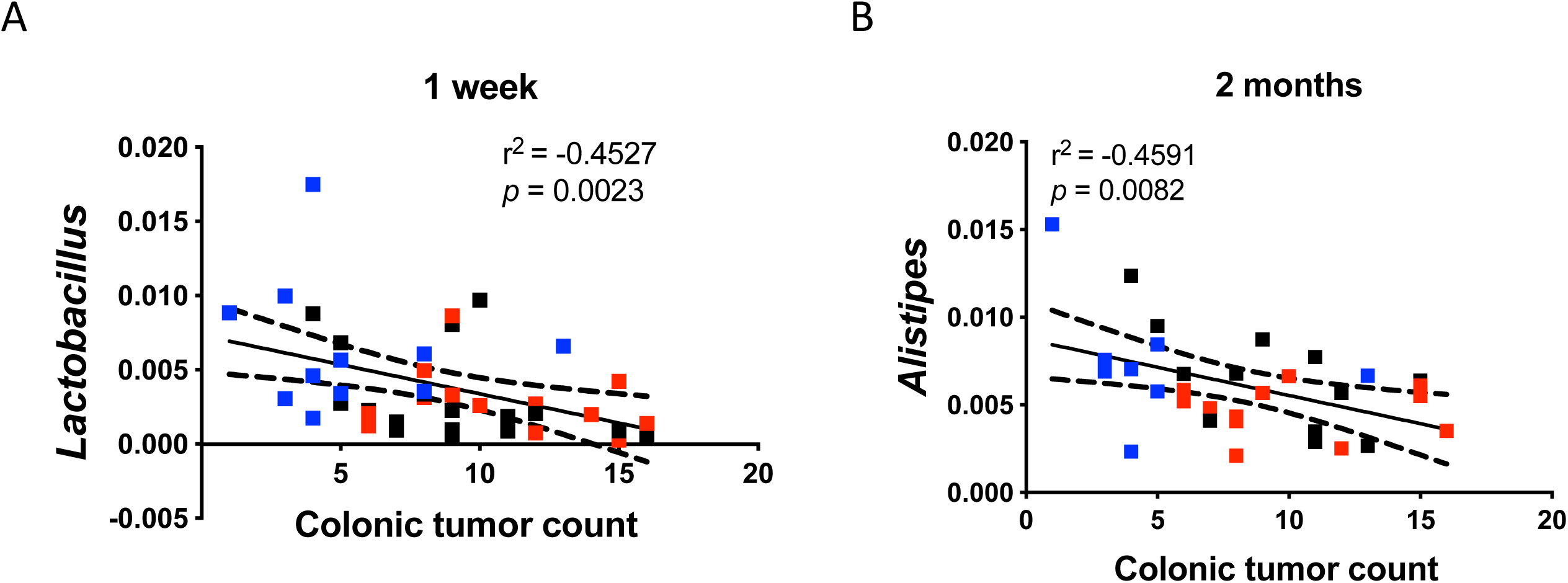
OTU-tumor correlations in control and treated PIRC rats. Pearson’s correlations (*p*<0.05) between OTU relative abundance from 2-month fecal samples with colonic tumor counts. Representative example of a negative correlation, *Lactobacillus* (A) and *Alistipes* (B) with colonic tumor count along x-axis and relative abundance of the taxa along the y-axis is shown.

## Supplementary Tables

**Supplementary Table 1:**
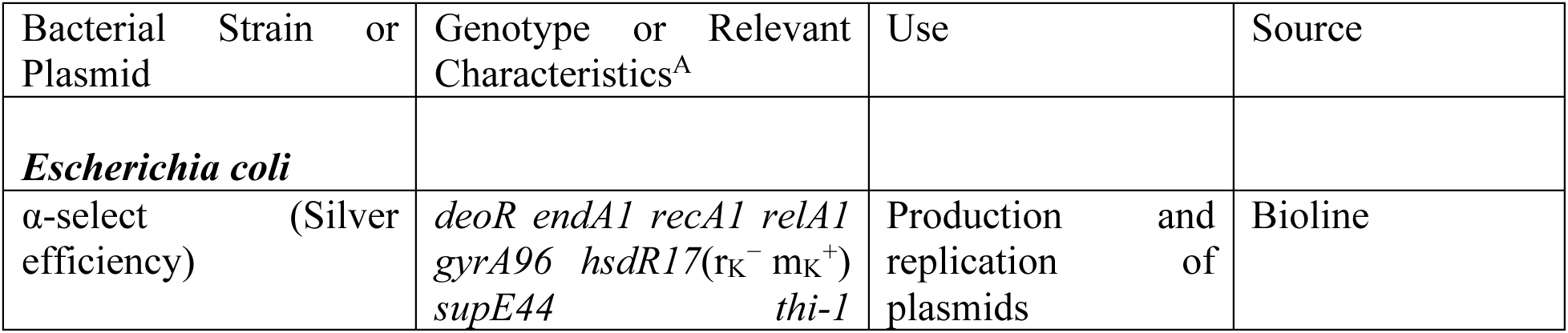
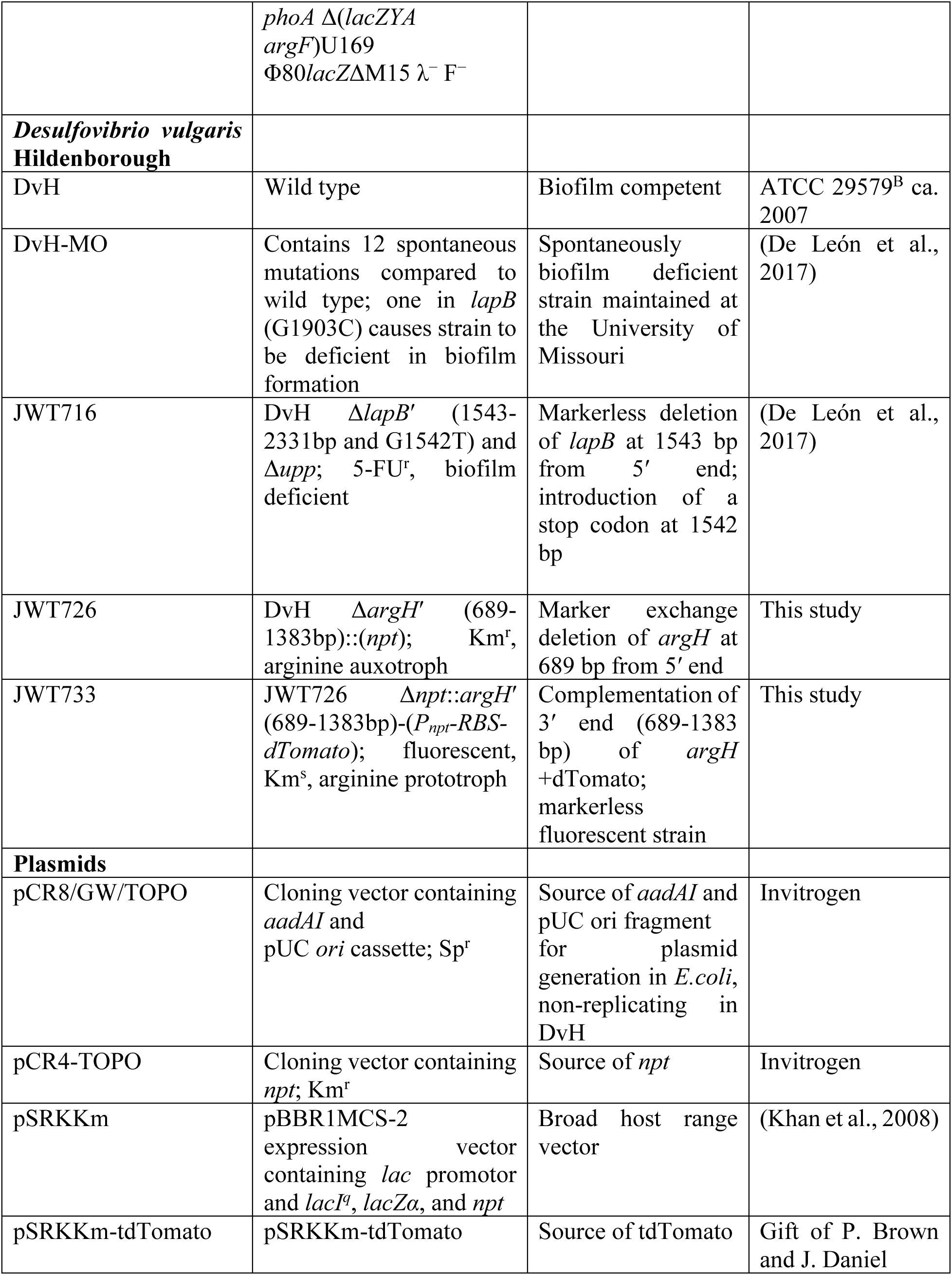
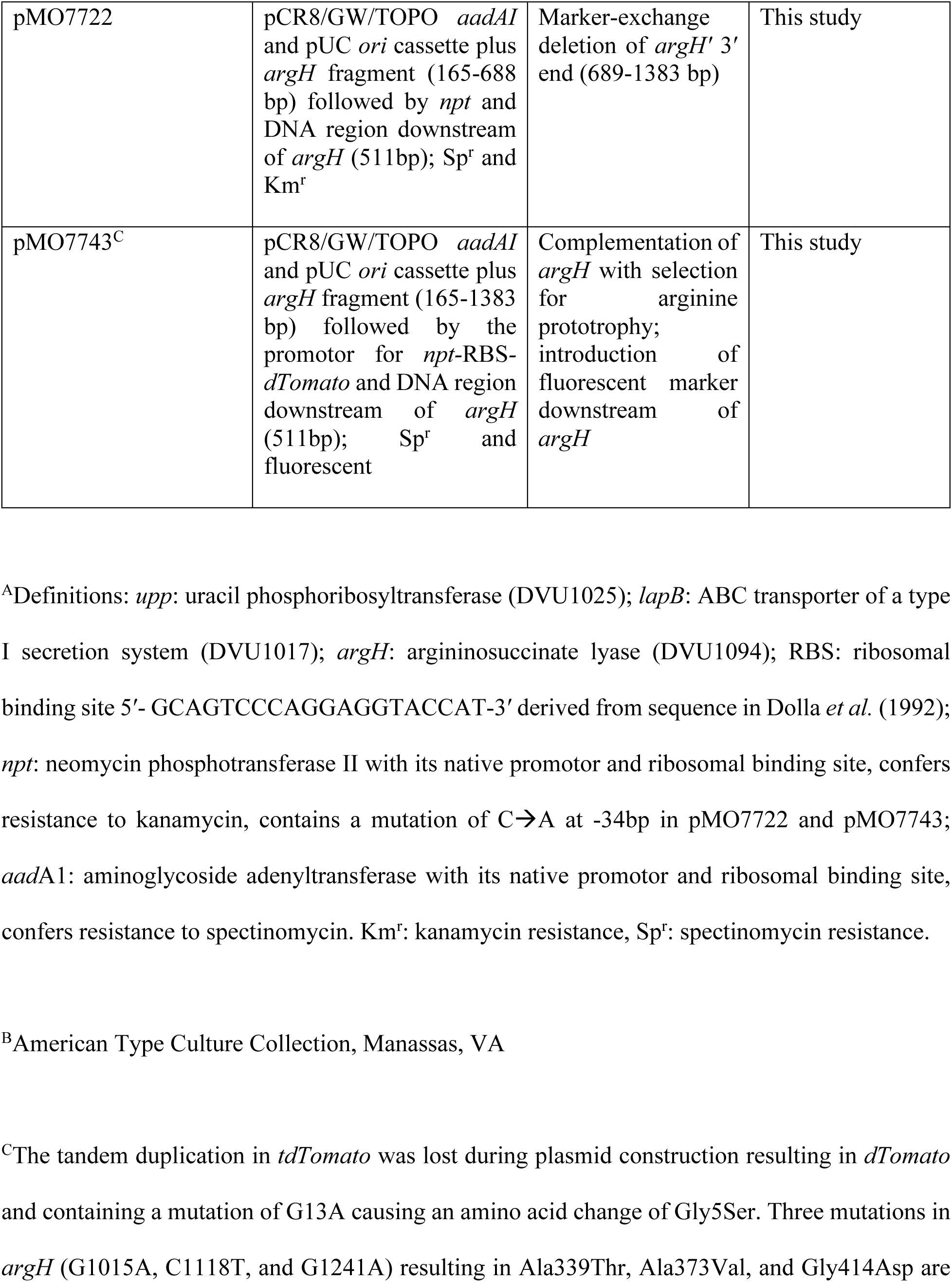
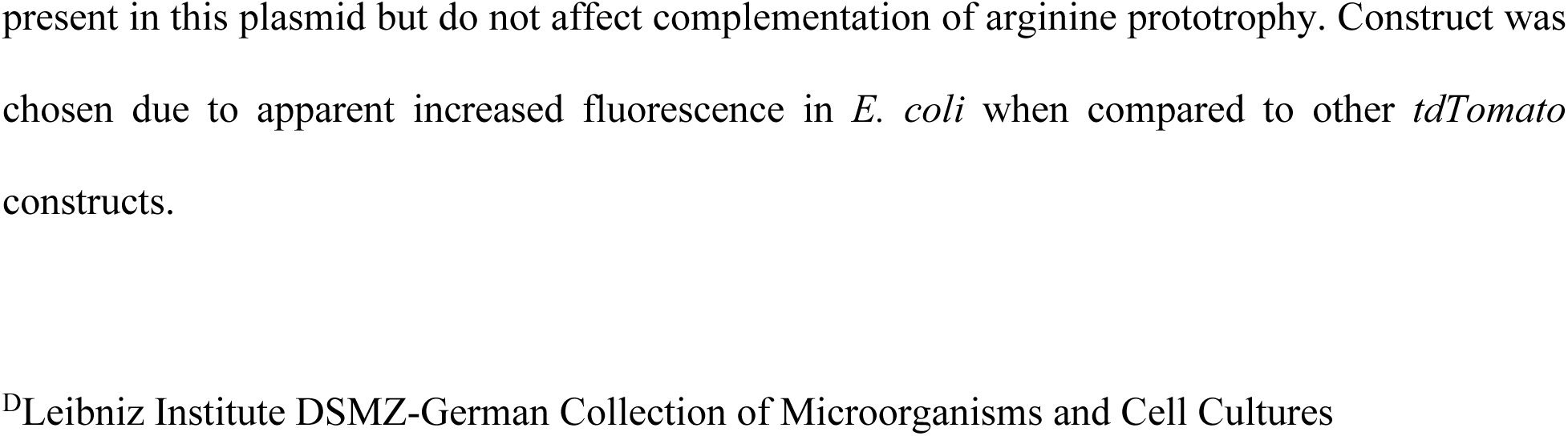
Bacterial strains and plasmids used in the study

**Supplementary Table 2:**
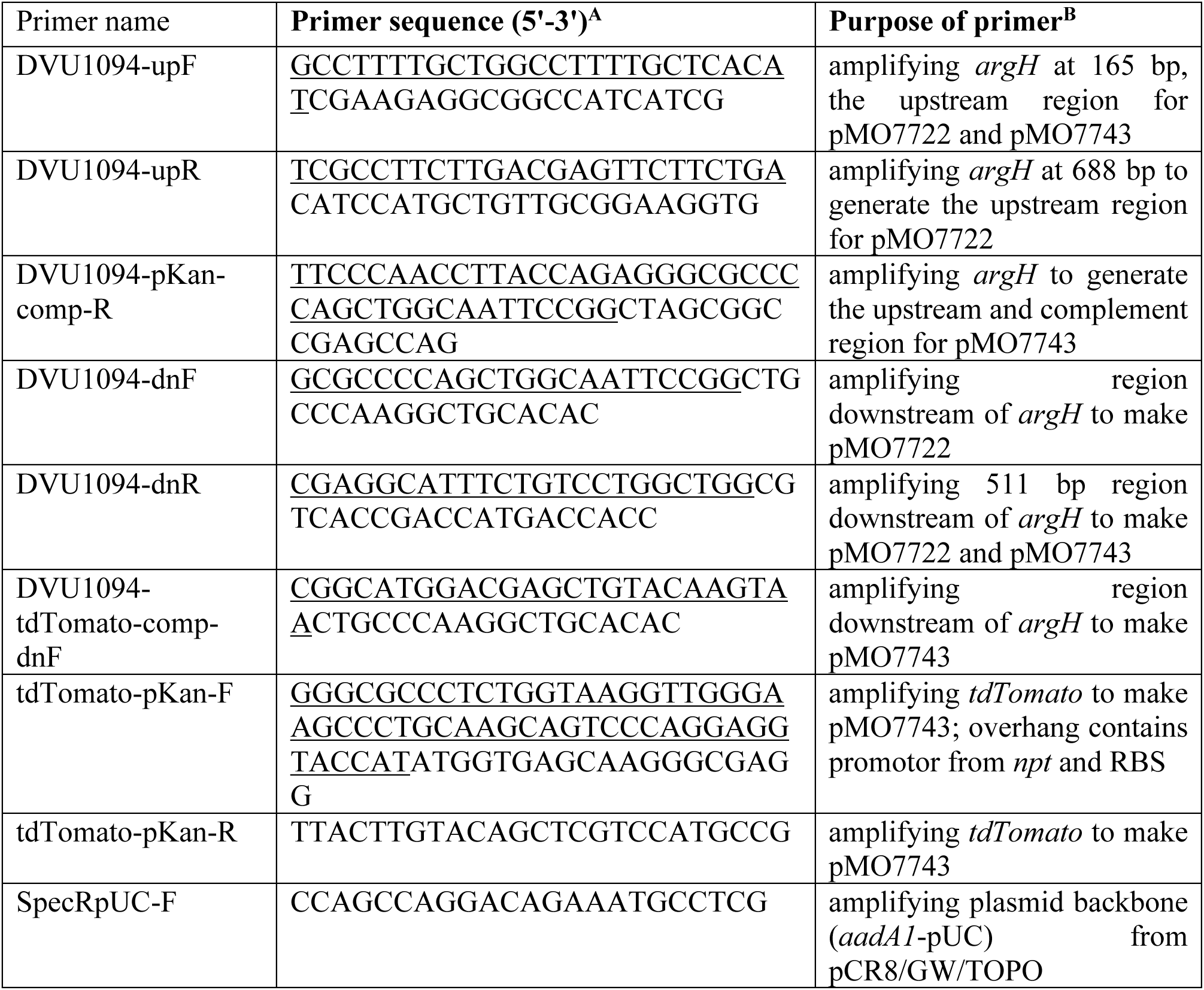
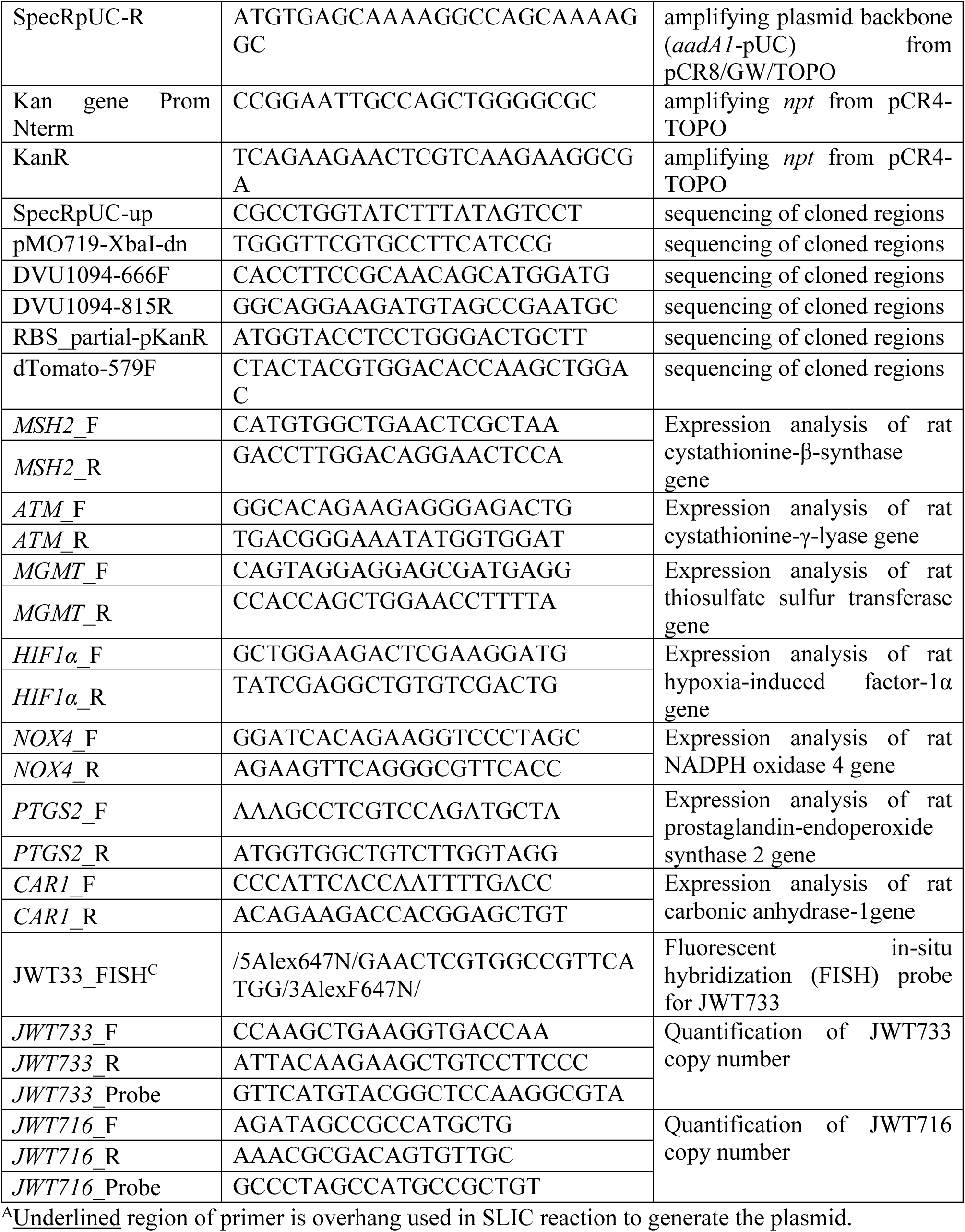
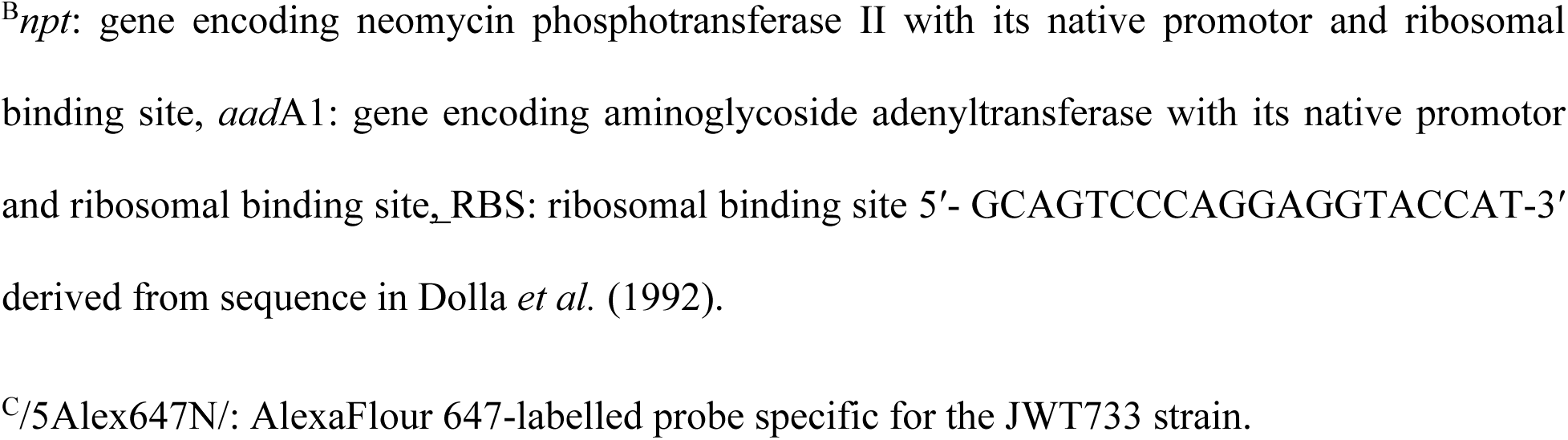
Primer and probes used in this study

**Supplementary Table 3:** Two-Way PERMANOVA post-hoc analysis of GM community profile in fecal and biopsy samples collected at 4 months of age

| Bonferroni-corrected *p* values |  |  |  |  |  |
| --- | --- | --- | --- | --- | --- |
| Timepoint/Group |  | 1 week |  | 4-mo |  |
|  |  | DvH | DvH-MO | DvH | DvH-MO |
| 1 week | DvH | 1 | 0.0158 | 0.0261 | 0.005 |
|  | DvH-MO | 0.0158 | 1 | 0.0079 | 0.0022 |
| 4-mo | DvH | 0.0261 | 0.0079 | 1 | 0.0147 |
|  | DvH-MO | 0.005 | 0.0022 | 0.0147 | 1 |

**Supplementary Table 4:** One-Way PERMANOVA post-hoc analysis of GM community profile in fecal samples from DvH-treated rats

| Bonferroni-corrected *p* values |  |  |  |  |
| --- | --- | --- | --- | --- |
|  | JWT733 | JWT716 | DvH | DvH-MO |
| JWT733 |  | 0.1162 | 0.1619 | 0.0109 |
| **JWT716** | 0.1162 |  | 0.3188 | **0.0222** |
| **DvH** | 0.1619 | 0.3188 |  | **0.0361** |
| **DvH-MO** | **0.0109** | **0.0222** | **0.0361** |  |

**Supplementary Table 5:** Two-Way PERMANOVA post-hoc analysis of GM community profile in fecal and biopsy samples collected at 4 months of age

| **Bonferroni-corrected *p* values** |  |  |  |  |  |  |
| --- | --- | --- | --- | --- | --- | --- |
| **Samples/Group** | **Fecal** |  |  | **Biopsy** |  |  |
|  | **PBS** | **JWT733** | **JWT716** | **Biopsy-PBS** | **Biopsy-JWT733** | **Biopsy-JWT716** |
| **PBS** |  | 1 | 0.063 | **0.0015** | **0.039** | **0.0015** |
| **JWT733** | 1 |  | 0.071 | **0.0015** | **0.0375** | **0.0015** |
| **JWT716** | 0.063 | 0.071 |  | **0.0015** | **0.0015** | **0.0015** |
| **Biopsy-PBS** | **0.0015** | **0.0015** | **0.0015** |  | 0.1815 | **0.0015** |
| **Biopsy-JWT733** | **0.039** | **0.0375** | **0.0015** | 0.1815 |  | 1 |
| **Biopsy-JWT716** | **0.0015** | **0.0015** | **0.0015** | **0.0015** | 1 |  |

## Notes

https://www.ncbi.nlm.nih.gov/bioproject/?term=PRJNA495020

### Competing Interest Statement

The authors have declared no competing interest.

